# Genome-wide association studies identify promising QTL for freezing tolerance in winter and early spring as a basis for in-depth genetic analysis and implementation in winter faba bean (*Vicia faba* L.) breeding

**DOI:** 10.1101/2024.11.25.625124

**Authors:** Alex Windhorst, Cathrine Kiel Skovbjerg, Deepti Angra, Donal Martin O’Sullivan, Stig Uggerhøj Andersen, Link Wolfgang

## Abstract

Interest in faba bean as a locally adapted high-protein grain legume crop has increased in Europe over the past decade. Winter faba bean, which can make use of soil moisture from autumn to spring and partially escape summer droughts, exhibit greater yield potential than the spring-type. However, due to insufficient winter hardiness, winterkill is a major constraint that prevents large-scale production of current winter-type cultivars in Central and Northern Europe. Here, we extend the understanding of freezing tolerance, the main component trait of winter hardiness, during winter and against late-frost in early spring and define genomic target regions for marker-assisted selection. Comparative analysis of genome-wide association studies revealed 13 treatment-specific major QTLs with partially pleiotropic effect on four freezing tolerance related traits. In addition, we identified five treatment-unspecific pleiotropic QTLs, including two major freezing tolerance loci on chromosomes 1 and 5. Our results thus indicate both a distinct and common genetic control of tolerance to winter- and late-frost in winter faba bean. In combination with the promising prediction abilities obtained from marker score-based prediction, our work highlights the potential for marker-assisted and genomic selection toward improved freezing tolerance and winter hardiness in winter faba bean breeding programs.

## Introduction

Winter faba bean can yield substantially higher (<47%, Link et al. 2010) than spring-type faba bean. Current winter faba bean production, however, is restricted to the UK and southern coastal regions of France (Link et al. 2010; Sass 2022), where the maritime climate favors mild winters. The large-scale production in the cooler regions of Central and Northern Europe, such as Poland, Denmark, and Germany, is limited by the risk of winter kill driven by an insufficient winter hardiness in current winter-type cultivars; spring faba bean dominates the market.

No two winters are alike and each challenges a winter crop with a broad palette of abiotic and biotic stresses, such as severe freezing, freeze-thaw cycles, hail, snow coverage, and snow mold. Thus, winter hardiness is a complex highly quantitative trait strongly influenced by the environment. Freezing tolerance (FT) has been reported as major component trait of winter hardiness (Pulli et al. 1996). The physiological process of acquirer FT is induced by shorter day length (Beck et al. 2004; Junttila 1996) alongside low, non-freezing temperatures and is referred to as cold acclimation or hardening (Thomashow 1999). Key regulon gene families such as the CBF/DBEF (Thomashow 1999) and multiple, partially independent signaling pathways have been identified to initiate hardening (Thomashow 1999; Xin and Browse 2000). Expression of cold responsive genes leads to the accumulation of sucrose, proline, and other cryoprotectants as well as modifications in membrane lipid composition to prevent frost injuries (reviewed by Xin and Browse 2000). Non-hardened plant tissue first shows symptoms of freeze-induced cellular dehydration (Thomashow 1999); phenotypically expressed as loss of turgor in leaves and stems. Severe frost can further lead to irreversible cell damage and cell membrane destruction, i.e. tissue blackening, and can be lethal (Thomashow 1999). The wide genetic variation for inherent FT prior to hardening and the level of FT acquired through it illustrates the polygenic complexity of this trait (Duc and Petitjean 1995; Harrison et al. 1997).

In spring, increasing day length and rising temperatures initiate the dehardening (Rognli 2013), i.e. reversion of cold acclimation and onset of plant growth. The dehardening rate depends on the temperature in spring. Cold nights can prevent dehardening even if day temperatures rise >10°C (Eagles and Williams 1992). Varying capacity for dehardening and quick re-hardening in response to frost spells during dehardening has been reported as well (Rognli 2013). In winter faba bean, dehardening starts at >7°C (Herzog 1989) and is likely enhanced by long-day conditions (Junttila 1996). Once dehardened, late-frost events in (early) spring are particularly harmful for the then-again frost sensitive plants (Duc et al. 2011; Maqpool et al. 2010).

Climate change leads to on average milder winters with an increase in unusual, severe early- and late-frost events (Francis and Skific 2015; Gu et al. 2008; Kalberer et al. 2006). To this end, sufficient hardening in autumn is at risk in winter crops. Milder fall temperatures have already been reported to affect hardening of pea and other grain legumes (Castel et al. 2017). Furthermore, mild winter conditions and fast temperature increase in early spring may provoke earlier and more rapid dehardening. Breeding toward improved winter hardiness and FT in winter and early spring is thus an important strategic goal to extend the area of cultivation of winter faba bean. Fortunately, promising genetic variation for FT in the genuine winter faba bean gene pool has been reported (Annicchiarico et al. 2015; Arbaoui et al. 2008; Carillo-Perdomo et al. 2022; Duc et al. 2011; Herzog 1987; Stoddard et al. 2006). However, breeding progress is limited, since assessment of winter hardiness in the field depends on rarely occurring well-differentiating cold winters. Instead, artificial screening under controlled conditions (Ali et al. 2016; Arbaoui et al. 2008; Duc and Petitjean 1995) allow for selection for FT (Annicchiarico and Iannucci 2007). However, correlation of FT screening results and winter survival in the field can be low (Waldron et al. 1998). Nevertheless, FT appears to be a highly heritable trait with large additive effect QTLs in winter faba bean (Ali et al. 2016; Carillo-Perdomo et al. 2022; Duc and Petitjean 1995), which offers promising potential for marker-assisted selection (MAS) and trait introgression. Hitherto, QTL-mapping and genome-wide association studies (GWAS) have identified several QTLs related to FT in winter faba bean (Ali et al. 2016; Arbaoui et al. 2008; Carillo-Perdomo et al. 2022; Sallam 2016). However, the available genomic tools utilized in previous studies did allow neither for fine mapping of QTLs nor for the identification of SNP markers directly applicable in MAS. Furthermore, no study has yet been published on the genetics of dehardening and late-frost tolerance in winter faba bean.

Due to recent achievements in developing faba bean genomic tools, such as 60K SNP Array (O’Sullivan et al. 2019), the Hedin/2 v.1 reference genome sequence (Jajakodi et al. 2023), and gene annotation browser (Jajakodi et al. 2023), more detailed genomic analyses of tolerance to freezing in winter (winter-frost; WF) and spring (late-frost; LF) became within reach. Our study therefore served a dual purpose: First, we aimed to further unravel the quantitative genetic architecture of the WF and LF tolerance. Therefore, we used phenotypic data of a n = 185 winter faba bean inbred line panel from independent artificial WF (Sallam 2014) and LF treatments (Windhorst et al. 2023). The two data sets provided the opportunity for a deep, comparative analysis under the assumption of an independent genetic control FT in hardened (WF) and dehardened (LF) winter faba bean. Our second objective was the GWAS-based identification of markers applicable in MAS for FT in winter faba bean breeding programs.

## Materials and Methods

### Plant Material

The Association-set (A-set; **Supplementary Table S.M.1**) and a faba bean Check-set (C-set) were utilized in this study. The origin and structure of the A-set (n = 188 winter faba bean inbred lines) were described elsewhere (cf. Arbaoui and Link 2008; Sallam 2014; Ali et al. 2016; Windhorst et al. 2023). The C-set comprised lines developed from in-house crosses and (old) cultivars (Windhorst et al. 2023): 24 winter-types (partially related to A-set) and four spring-types. The C-set was included as checks throughout all experiments.

### Phenotyping

The A-set was phenotyped for tolerance to artificial winter-frost (WF; cf. Sallam 2014) and late-frost (LF; cf. Windhorst et al. 2023) stress treatments in repeated climate chamber experiments (**Table 1**). Each experiment had two replicates randomized by α-lattice design with incomplete blocks. Juvenile potted plants were tested employing treatment-specific protocols. The WF protocol consisted of five steps: (1) germination and growth (30 days), (2) hardening (10 d), (3) frost test (4 d), (4) regeneration after frost (4 d), and (5) regrowth (30 d). The frost test was a standardized freeze-thaw cycle with three successive frost nights at -16, -18, and -19 °C. The WF protocol was adapted to test for LFT by adding two steps to the protocol between the hardening and the frost test: (1) the ‘winter’ (10 d), followed by (2) dehardening (10 d). Hence, dehardened plants were tested in the frost test at slightly increased test temperatures: -13, -15, and -17 °C.

**Table 1:** Artificial freezing tolerance experiments conducted in climate chamber.

| Treatment | Exp. Series | Set | no. Lines | No. of Exp. | Repl. per Exp. | Season (Sep-Apr) |
| --- | --- | --- | --- | --- | --- | --- |
| winter-frost (WF) | WFT | A-set | 185 | 10 | 2 | 2011-12, 2012-13 |
| late-frost (LF) | LFT | A-set | 183 | 5 | 2 | 2020-2021, 2021-22 |

Exp., experiment Repl., replication

We included three different categories of traits (**Table 2**): (1) morphology and growth-habit traits prior to freezing, also referred to as non-freezing-related sub-traits (non-FRS-traits), (2) frost stress symptom FRS-traits, and (3) survival-related FRS-traits. Plant height (PH), leaf number (LN), and number of tillers (Til) were assessed twice; after both hardening and dehardening (indicated by “1” and “2”). DeltaPH denotes the plant height increase during dehardening in the LF treatment. The loss of turgor and loss of color in leaves (and stem) were each visually scored after each frost night (T1, C1, T2, C2, T3, C3) on treatment specific scales (WF: 1-9; LF: 1-4; no to full symptoms) and summed across frost nights to total loss of turgor (LossT), total loss of color (LossC), and the total loss of turgor and color (LossTC). Disposition to survive (DtS; 0-90°) was the transformed days of survival after frost test of each genotype calculated after Roth and Link (2010). The shoot fresh matter weight (REG; g) of the surviving plants was measured at the end of the regrowth phase. Experiment-wise, REG was linearly transformed (T-REG) to a fixed maximum of 5 g (WF) or 10 g (LF) by taking the mean REG of the top five genotypes as transformation factor, respectively. Transformation was conducted to account for large between-experiment differences in REG (mean and variance). Unless stated differently, the term ‘trait’ always refers to all traits examined, including the FRS-traits.

**Table 2:** Traits and freezing-related sub-traits assessed in winter-frost and late-frost experiments.

| Trait-Type | Trait ID | Trait name | Unit | Scale | Treatment |  |
| --- | --- | --- | --- | --- | --- | --- |
|  |  |  |  |  | WF | LF |
| non-FRS-traits | PH1 | Plant height after hardening | cm |  | x | x |
|  | LN1 | Leaf number after hardening | count |  | x | x |
|  | Til1 | Number of tillers after hardening | count |  |  | x |
|  | PH2 | Plant height after dehardening | cm |  |  | x |
|  | LN2 | Leaf number after dehardening | count |  |  | x |
|  | Til2 | Number of tillers after dehardening | count |  |  | x |
|  | deltaPH | Change in plant height from PH1 to PH2 | cm |  |  | x |
| Symptom FRS-traits | T1 | Loss of turgor after 1st night of frost stress | score | 1-9; 1-4 | x | x |
|  | C1 | Loss of color after 1st night of frost stress | score | 1-9; 1-4 | x | x |
|  | T2 | Loss of turgor after 2nd night of frost stress | score | 1-9; 1-4 | x | x |
|  | C2 | Loss of color after 2nd night of frost stress | score | 1-9; 1-4 | x | x |
|  | T3 | Loss of turgor after 3rd night of frost stress | score | 1-9; 1-4 | x | x |
|  | C3 | Loss of color after 3rd night of frost stress | score | 1-9; 1-4 | x | x |
|  | LossT | Sum of turgor loss | score | 4-36; 8-32 | x | x |
|  | LossC | Sum of color loss | score | 4-36; 8-32 | x | x |
|  | LossTC | Sum of turgor and color loss | score | 8-72; 16-64 | x | x |
| Survival FRS-traits | DtS | Disposition to survive | grade [°] | 0-90 | x | x |
|  | T-REG | Transformed regrowth | weight [g] |  | x | x |

FRS-traits, freezing-related sub-traits

### Phenotypic Data Analysis

First, each experiment was analyzed according to the α-lattice design in PLABSTAT (Utz 2011). Thereafter, analysis of variance (ANOVA) was performed across experiments within each treatment using a mixed linear model in PLABSTAT. The **model 1** was fitted for the WFT (Ali et al. 2016) series:

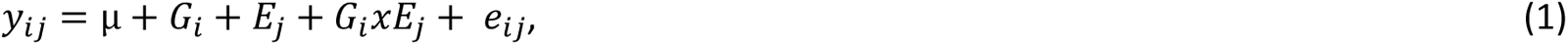

where *Y_ij_* denotes the lattice-adjusted mean of the *i*-th genotype in the *j*-th experiment (lattice), μ is the overall trait mean, *G_i_* is the random genetic effect of the *i*-th genotype, *E*j is the fixed effect of the *j*-th experiment, *G_i_xE_j_* denotes the random effect of the genotype-by-experiment interaction of the *i*-th genotype in the *j*-th experiment, and *e_ij_* is the residual error. Due to sometimes substantial differences in freezing injury levels observed between replicates of the same LFT experiment, lattice-adjusted values of the 10 single replicates (*R*) of the five LFT experiments were used instead of lattice-adjusted means per experiment to fit the **model 2**. Thus, *G_i_xE_j_* was not separated and the residual error *e_ij_* consisted of the *G_i_xR_k_* interaction effects:

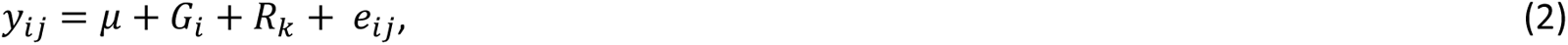

For WFT repeatability (h^2^) was calculated as:

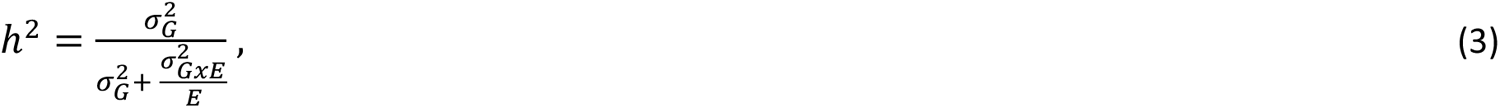

where 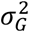 is the genotypic variance estimated in **model 1**, 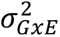 denotes the *GxE* variance (**Eq. 1**), and *E* is the number of experiments. For LFT we used:

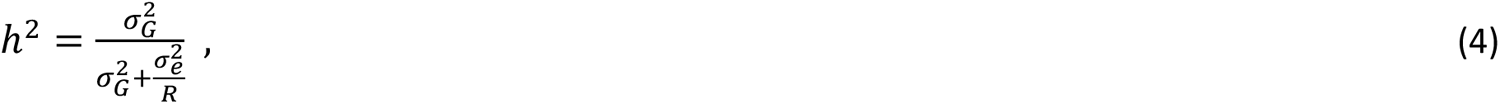

where 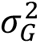 was estimated in **model 2**, *R* denotes the number of experiments and 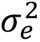 indicates the error variance. Results from **models 1** and **2** were utilized in GWAS and to calculate phenotypic Pearson correlation for all trait pairs within and across treatments. Genotypic correlations were calculated between all traits within treatments.

### SNP Genotyping

The A-set was genotyped with the Affymetrix 60k Vfaba_v2 Axiom SNP array (O’Sullivan 2019). DNA extraction, quality control, initial filtering of the genotyping data, and alignment of high-quality SNPs to the *V. faba* Hedin/2 v.1 reference genome sequence (Jayakodi et al. 2023) was described in Skovbjerg et al. (2023). In the following, chromosome 1 will be displayed as chr1S and chr1L; split at position 1,574,527,093 (Skovbjerg et al. 2023). We performed pairwise redundancy check between genotyped samples by calculating genetic identities (GI) based on 21,345 high-quality SNPs following Skovbjerg et al. (2023):

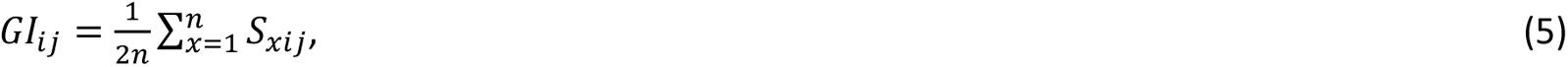

where *GI_ij_* is the GI between the *i*-th and *j*-th genotype sample, *n* is the number of SNPs where neither of the two samples shows missingness, and *S_xij_* is the number of shared alleles between samples *i* and *j* at SNP *x*. *S_xij_* could take values of 0, 1, and 2. In case two samples had a GI ≥90%, one was discarded. The GI threshold was set based on duplicated check genotyping samples. After filtering for line heterozygosity (<5%), 185 and 183 A-set lines remained in the WF and LF dataset. Within these treatment specific A-set versions, we removed monomorphic SNPs and applied filters for SNP heterozygosity (≤0.10), SNP missingness (≤0.10), and minor allele count (≥10). Final SNP-sets sizes were 17,227 SNPs for the WF A-set and 17,211 SNPs for the LF A-set, with a share of 17,205 SNPs. Missing SNP information was imputed with the LD-kNNi algorithm on default settings in TASSEL 0.5 (Bradbury et al. 2007). The SNP density was calculated in relation to physical and genetic map positions (NV664xNV153; Björnsdotter et al. 2022) and displayed with CMplot (v.4.5.0; LiLin 2023) R package. Principal component analysis (PCA) was performed with 19,791 polymorphic SNPs, prior to filtering, in the A-set (n = 185) using adegenet (v.2.1.4; Jombart and Ahmed 2011) and ade4 (v.1.7-22; Dray and Dufour 2007) in R environment (v.4.0.3; R Core Team 2021).

### Genome-Wide Association Study

We performed GWAS using the Bayesian-information and Linkage-disequilibrium Iteratively Nested Keyway (BLINK; Huang et al. 2019) model at default settings (GAPIT v.3; Wang and Zhang 2021). Population structure was controlled through trait-optimized number of principal components (PCs; range: 1-8; **Supplementary Table S.M.2**) as covariates. The optimum PC number was deduced from quantile-quantile (QQ) plots by selecting for the best fitting p-values. This strategy was chosen due to the substantial impact of the number of PCs as covariates on the GWAS results has been reported (Malik et al. 2019; Meyer et al. 2021; Khodaeiaminjan et al. 2023). We accounted for relationship among lines by kinship matrix (VanRaden 2008). Significance of marker-trait associations (MTAs) was tested by false discovery rate (FDR; Benjamini and Hochberg 1995) at a threshold of *p*(FDR) ≤0.05. Marker effect sizes and phenotypic variance explained (PVE in %) are reported from BLINK output. QQ and Manhattan plots were created with CMplot.

### Pleiotropic QTL Identification

Linkage Disequilibrium (LD) was estimated in the n = 185 A-set. Pairwise LD (r^2^) for each pair of SNPs was computed in PLINK (v 1.9; Prucell et al. 2007) and LD heatmaps generated in R (LDheatmap v.1.0-5; Shin et al. 2006). LD decay, defined as the distance to which genome-wide or chromosome-wide LD decayed to half of its maximum, was calculated as described by Windhorst et al. (2023). Putative pleiotropic QTL (plQTL) were identified both within and across treatments: Two markers were considered to address the same putative plQTL if they were within twice the chromosome-wide LD decay distance.

### Marker Score

For each A-set line we calculated trait-specific marker scores 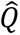 (Lande and Thompson 1989; Puspitasari et al. 2022; Windhorst et al. 2023) as:

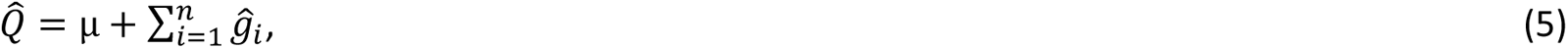

where µ is the A-set phenotypic mean, *n* equals the number of A-set lines, and 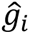 is the BLINK-estimated marker allele effect at the *i*-th marker locus. Note that per line only homozygous marker loci were considered. Prediction ability, i.e. Pearson correlation of 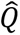 and phenotypic values, and explained variance (R^2^; squared correlation coefficient) were reported.

## Results

### WFT and LFT Phenotypic Data

Analysis of freezing tolerance phenotypic data from WF treatment in the A-set (WFT series) was reported in detail beforehand (Ali et al. 2016; Sallam 2014) and is briefly summarized here. In the ANOVAs, significant variation (p ≤0.01) due to genotype and experiment was found for all traits in the WFT and LFT series and for GxE interaction in WFT series also (**Table 3**). In LFT series, repeatability (h²) was high and ranged from 62.58-92.58%, with h² >70% in most FRS-traits. Similarly, high h² (>90%) were found for most traits in WFT series. Interestingly, h² was always slightly higher for color loss than for turgor loss in both series.

**Table 3:** Summary of phenotypic data and ANOVA of winter-frost and late-frost experiments with Association-set. All tested variances were significant (p ≤0.05).

| Trait | Series | Mean | Min | Max | CoV | VarG | VarE | VarGxE | VarError | h<sup>2</sup> |
| --- | --- | --- | --- | --- | --- | --- | --- | --- | --- | --- |
| PH1 | WFT | 5.80 | 4.06 | 7.95 | 0.15 | 0.68 | 0.35 | 0.14 | 0.47 | 90.93 |
|  | LFT | 3.35 | 1.62 | 6.95 | 0.23 | 5.13 | 26.10 | - | 6.07† | 89.41 |
| LN1 | WFT | 1.96 | 1.34 | 2.51 | 0.09 | 0.03 | 0.08 | 0.01 | 0.04 | 83.00 |
|  | LFT | 4.89 | 3.78 | 6.47 | 0.11 | 2.80 | 7.24 | - | 3.58† | 88.67 |
| Til1 | WFT | - | - | - | - | - | - | - | - | - |
|  | LFT | 0.11 | -0.07 | 0.61 | 1.11 | 0.10 | 0.20 | - | 0.60† | 62.58 |
| PH2 | WFT | - | - | - | - | - | - | - | - | - |
|  | LFT | 6.68 | 4.03 | 11.51 | 0.18 | 14.23 | 48.66 | - | 11.41† | 92.58 |
| LN2 | WFT | - | - | - | - | - | - | - | - | - |
|  | LFT | 9.14 | 6.97 | 12.23 | 0.11 | 8.24 | 32.77 | - | 16.21† | 83.56 |
| Til2 | WFT | - | - | - | - | - | - | - | - | - |
|  | LFT | 0.78 | 0.00 | 1.62 | 0.42 | 0.89 | 1.09 | - | 1.98† | 81.82 |
| delatPH | WFT | - | - | - | - | - | - | - | - | - |
|  | LFT | 3.33 | 2.14 | 5.47 | 0.20 | 3.85 | 18.36 | - | 5.25† | 88.01 |
| T1 | WFT | 3.33 | 1.92 | 4.91 | 0.17 | 0.29 | 0.27 | 0.08 | 0.35 | 85.78 |
|  | LFT | 5.80 | 4.26 | 7.23 | 0.10 | 2.34 | 6.76 | - | 9.72† | 70.67 |
| C1 | WFT | 2.50 | 1.58 | 3.67 | 0.17 | 0.14 | 0.15 | 0.01 | 0.37 | 76.80 |
|  | LFT | 4.19 | 3.05 | 5.94 | 0.13 | 2.15 | 11.14 | - | 6.96† | 75.50 |
| T2 | WFT | 4.45 | 2.50 | 6.27 | 0.17 | 0.49 | 0.20 | 0.09 | 0.45 | 89.09 |
|  | LFT | 6.43 | 4.93 | 7.96 | 0.09 | 2.49 | 5.24 | - | 8.86† | 73.72 |
| C2 | WFT | 4.00 | 2.25 | 6.26 | 0.20 | 0.60 | 0.22 | 0.08 | 0.49 | 90.51 |
|  | LFT | 5.25 | 3.77 | 6.83 | 0.10 | 2.41 | 9.31 | - | 6.73† | 78.17 |
| T3 | WFT | 5.33 | 3.17 | 7.23 | 0.16 | 0.63 | 0.33 | 0.13 | 0.53 | 89.59 |
|  | LFT | 6.71 | 5.28 | 7.91 | 0.08 | 2.08 | 4.01 | - | 8.88† | 70.11 |
| C3 | WFT | 4.93 | 3.12 | 7.21 | 0.18 | 0.72 | 0.41 | 0.09 | 0.55 | 91.07 |
|  | LFT | 5.77 | 4.47 | 7.11 | 0.09 | 2.19 | 7.71 | - | 6.16† | 78.04 |
| LossT | WFT | 18.55 | 11.01 | 25.72 | 0.16 | 7.95 | 3.40 | 1.61 | 5.68 | 90.76 |
|  | LFT | 18.93 | 15.17 | 22.85 | 0.08 | 20.17 | 45.51 | - | 61.23† | 76.71 |
| LossC | WFT | 16.73 | 10.54 | 24.47 | 0.17 | 7.87 | 3.81 | 0.96 | 5.13 | 92.09 |
|  | LFT | 15.21 | 11.49 | 19.78 | 0.10 | 19.47 | 79.31 | - | 40.31† | 82.85 |
| LossTC | WFT | 35.28 | 21.96 | 50.21 | 0.16 | 30.25 | 12.18 | 4.76 | 19.48 | 91.83 |
|  | LFT | 34.14 | 28.11 | 42.48 | 0.08 | 68.66 | 230.48 | - | 159.18† | 81.18 |
| DtS | WFT | 65.71 | 40.44 | 84.57 | 0.15 | 84.88 | 22.64 | 28.11 | 142.06 | 79.12 |
|  | LFT | 63.62 | 24.62 | 89.71 | 0.20 | 1263.88 | 1395.25 | - | 2882.45† | 81.43 |
| T-REG | WFT | 1.11 | 0.15 | 2.84 | 0.55 | 0.25 | 0.12 | - | 1.14† | 68.58 |
|  | LFT | 2.20 | -0.08 | 4.99 | 0.49 | 8.13 | 7.21 | - | 51.86† | 70.17 |

† No significance test conducted
Cov, Coefficient of variation; VarG, genotypic variance; VarE, experimental variance; VarGxE, genotype-by-experiment variance; VarError, error variance; h<sup>2</sup>, repeatability

We found significant, moderate to high phenotypic and genetic correlations between most traits within both series (**Supplementary Tables S.R.1-2**). DtS and T-REG were each strongly negatively correlated with all turgor and color loss traits. PH2 and deltaPH correlated stronger with all FRS-traits than PH1 in LFT.

Interestingly, both extent and patterns of correlation of PH1 (LF) with all WF traits were very similar to the correlations observed for PH1 (WF) with all WF traits (**Supplementary Table S.R.3**). Except for LN1, patterns of correlations were symmetrically among the remaining traits between the treatments (WFT vs LFT).

### SNP Density and PCA

The 17,227 SNPs and 17,211 SNPs that remained in the WFT SNP-set and the LFT SNP-set after quality control covered all chromosomes and the entire genome quite well (**Fig. 1**). The average SNP density was 1.52 SNPs/Mb on the physical and 8.85 SNP/cM on the genetic map (Supplementary Table S.R.7). The average SNP-to-SNP distance was 0.67 Mb or 0.11 cM and the average SNP-to-mid-gene distance was 0.14 Mb at an average SNP/gene ratio of 0.49 (∼ 1 SNP/2 genes; **Supplementary Table S.R.4**). In the A-set, LD decayed genome-wide after 1.32 Mb to half of its maximum but considerably slower chromosome-wide (17.57 Mb; **Supplementary Table S.R.5**). The LD observed at the average SNP-to-SNP distance (Mb) was r² = 0.15, while the average LD was r² = 0.05 (**Supplementary Table S.R.4**). As expected, PCA showed no substantial structuring within the A-set (**Supplementary Figure S.R.1**).

**Fig. 1:**
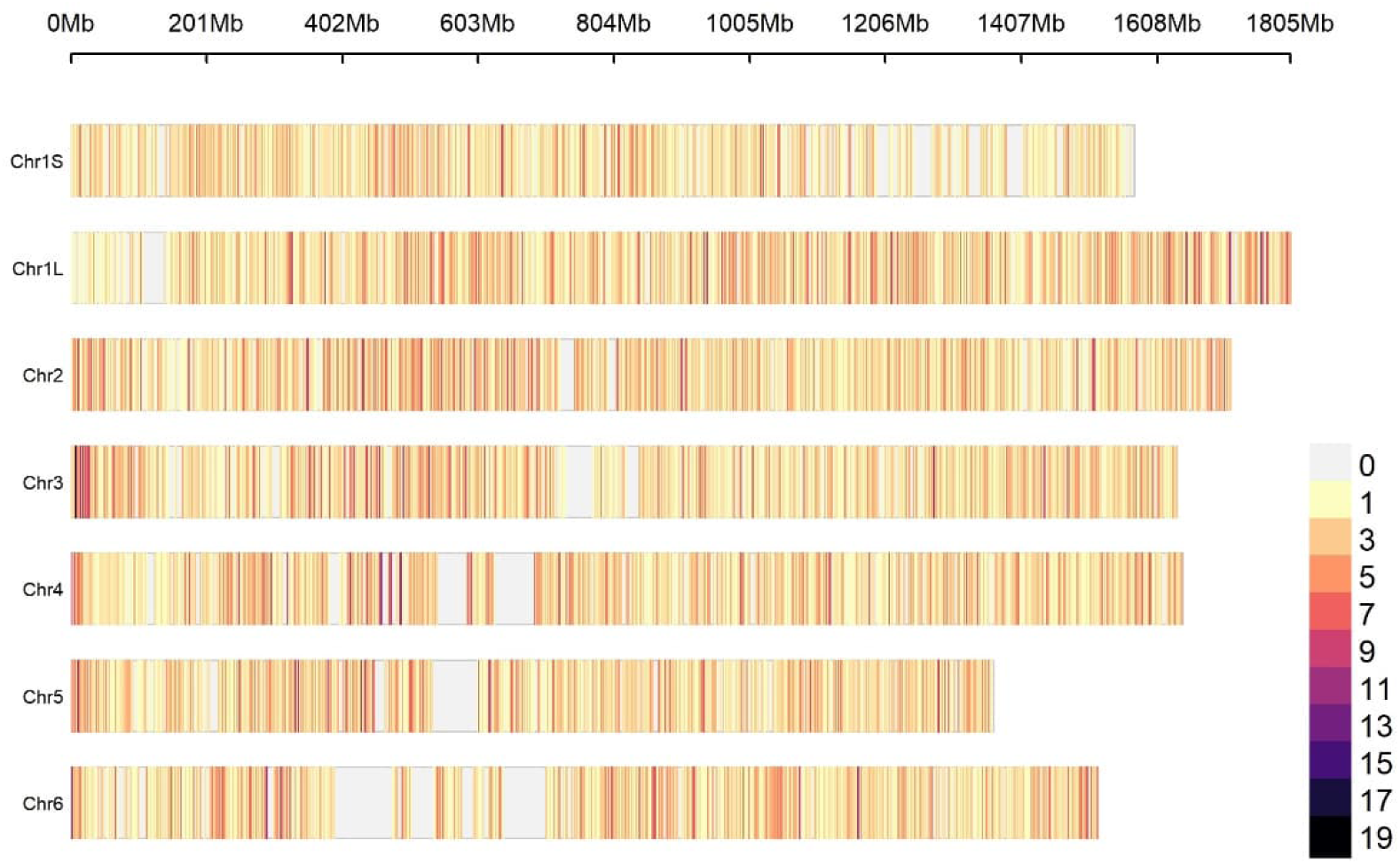
SNP density across faba bean chromosomes based on the 17,227 SNPs in the WF SNP-set. Color code: SNP density in 1 Mb window size.

### GWAS on WFT

The GWAS carried out for the 13 WF traits yielded at least two significant MTAs per trait and 57 in total (**Fig. 2**; **Table 4**; **Supplementary Figure S.R.2**). Of these, 37 MTAs were based on unique markers while the remaining 20 MTAs were based on ten markers. The marker AX-416774176, for instance, was significantly associated with all turgor and color loss FRS-traits, except C3, LossC, and LossTC. Two more markers (AX-416766819, AX-416775062) were associated with four turgor and color loss traits each. Only one marker (AX-181158344) was significant for two different trait categories: the survival FRS-traits DtS and the symptom FRS-traits T1 and T3. We found the most MTAs for T3 (n = 8) followed by C1, LossC, and DtS with six MTAs each. Single marker PVE ranged from 0.99% (T3; AX-416776564) to 32.90% (C2, AX-416774176). The trait-wise total PVE by all markers ranged from 25.07% (T-REG) to 58.12% (T3; **Table 6**). Among the 37 unique markers, often the major allele (n = 23) had the beneficial effect on trait expression.

**Fig. 2:**
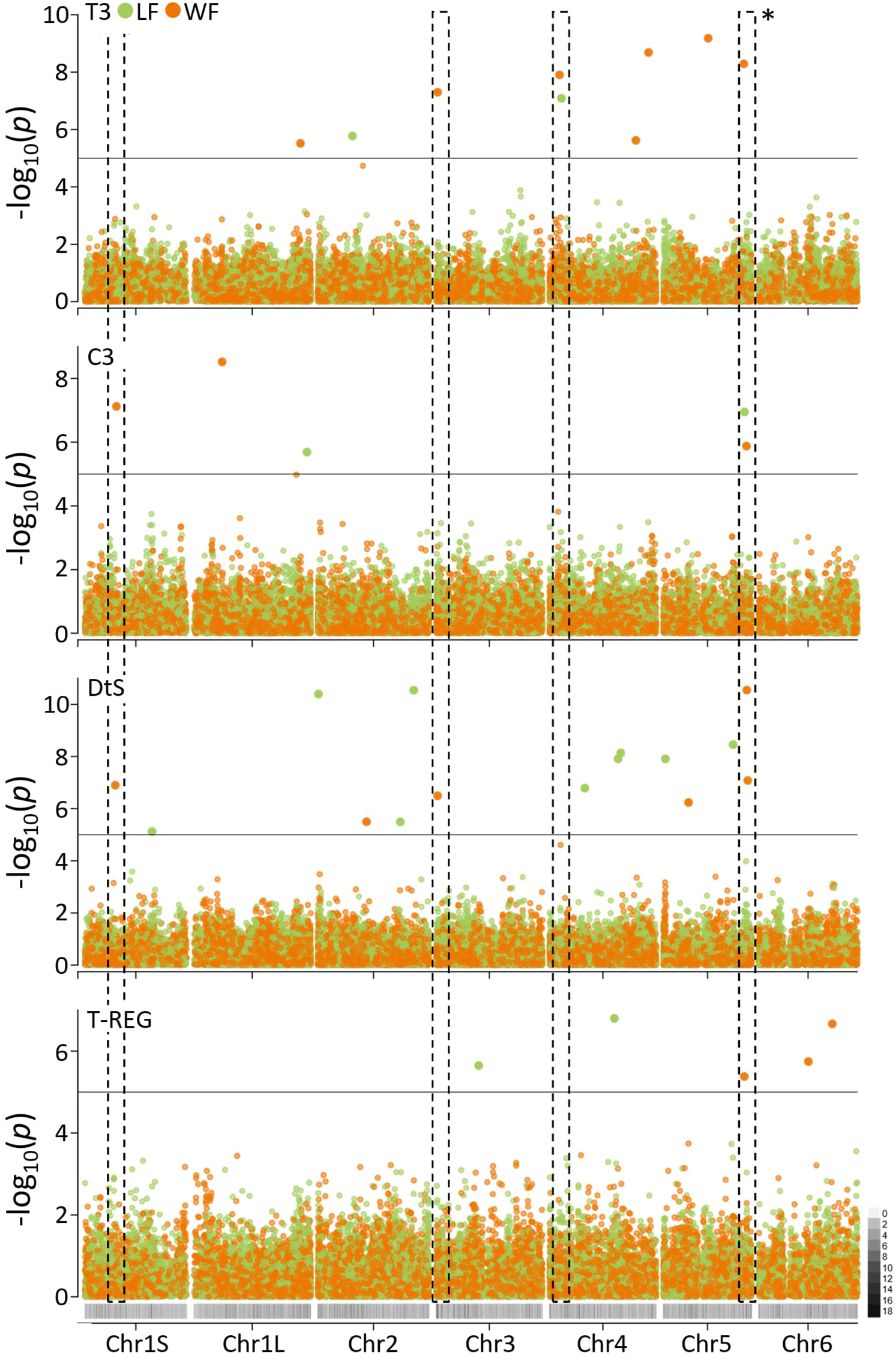
Manhattan plot of genome-wide association analysis results for traits related to freezing tolerance in winter-frost (orange) and late-frost (green) treatment. Top to bottom: Turgor (T3) and (C3) color loss (C3) both after third frost stress during frost treatment; disposition to survive (DtS) after frost treatment; transformed regrowth of survivors (T-REG). Grey color code: SNP density along chromosome (Fig. 1). **\***, highlighted pleiotropic QTL starring in Fig. 3.

**Table 4:** Catalog of significant marker-trait associations from genome-wide association study for winter-frost tolerance. In descending order per trait according to p-value.

| Trait | SNP_ID | Chr | Pos Mb | ben-Allele | ben-AF | ben-Ef-fect | PVE % | plQTL | MTA in LF |
| --- | --- | --- | --- | --- | --- | --- | --- | --- | --- |
| PH1 | AX-416801956 | 4 | 1571.44 | A | 0.86 | -0.43 | 17.72 |  | PH2 |
| PH1 | AX-181464412 | 5 | 225.69 | G | 0.75 | -0.29 | 7.42 |  |  |
| PH1 | AX-181440894 | 1L | 1225.51 | G | 0.49 | -0.23 | 4.53 |  |  |
| LN1 | AX-416801758 | 4 | 1135.55 | T | 0.91 | -0.11 | 22.21 |  |  |
| LN1 | AX-416776892 | 6 | 1050.05 | G | 0.66 | -0.06 | 5.27 |  |  |
| LN1 | AX-181176222 | 3 | 311.88 | C | 0.74 | -0.06 | 4.83 |  |  |
| LN1 | AX-416746325 | 4 | 381.86 | A | 0.22 | -0.06 | 6.85 |  |  |
| T1 | AX-416774176<sup>a</sup> | 5 | 693.09 | C | 0.94 | -0.42 | 16.97 | WFc5PI-1 |  |
| T1 | AX-416766819<sup>b</sup> | 4 | 1523.15 | T | 0.12 | -0.29 | 5.81 | WFc4PI-2 |  |
| T1 | AX-181158344<sup>c</sup> | 3 | 16.66 | C | 0.96 | -0.42 | 16.58 | WFc3PI-1 | T2, LossT |
| T1 | AX-181494082<sup>d</sup> | 4 | 146.92 | G | 0.96 | -0.42 | 16.75 | WFc4PI-1 |  |
| C1 | AX-416774176<sup>a</sup> | 5 | 693.09 | C | 0.94 | -0.35 | 21.08 | WFc5PI-1 |  |
| C1 | AX-416775266 | 3 | 647.95 | G | 0.86 | -0.17 | 5.57 |  |  |
| C1 | AX-181199308 | 4 | 1544.72 | G | 0.05 | -0.27 | 7.14 | WFc4PI-2 |  |
| C1 | AX-416815923 | 3 | 1230.49 | G | 0.41 | -0.12 | 2.12 |  |  |
| C1 | AX-181491337 | 1L | 905.23 | A | 0.89 | -0.18 | 7.28 |  |  |
| C1 | AX-416813597 | 1S | 1044.45 | G | 0.65 | -0.10 | 1.95 |  |  |
| T2 | AX-416775062<sup>e</sup> | 2 | 694.64 | C | 0.45 | -0.24 | 4.93 |  |  |
| T2 | AX-416789006<sup>f</sup> | 6 | 1134.78 | C | 0.38 | -0.22 | 3.27 | WFc6PI-1 |  |
| T2 | AX-416799928<sup>g</sup> | 4 | 1413.44 | C | 0.88 | -0.27 | 4.96 |  |  |
| T2 | AX-416774176<sup>a</sup> | 5 | 693.09 | C | 0.94 | -0.39 | 17.33 | WFc5PI-1 |  |
| C2 | AX-416766819<sup>b</sup> | 4 | 1523.15 | T | 0.12 | -0.38 | 10.05 | WFc4PI-2 |  |
| C2 | AX-416774176<sup>a</sup> | 5 | 693.09 | C | 0.94 | -0.52 | 32.90 | WFc5PI-1 |  |
| C2 | AX-416724438<sup>h</sup> | 1S | 483.98 | G | 0.80 | -0.28 | 6.06 | WFc1SPI-1 |  |
| T3 | AX-416774176<sup>a</sup> | 5 | 693.09 | C | 0.94 | -0.53 | 13.76 | WFc5PI-1 |  |
| T3 | AX-416766819<sup>b</sup> | 4 | 1523.15 | T | 0.12 | -0.37 | 4.59 | WFc4PI-2 |  |
| T3 | AX-416798162 | 5 | 1238.85 | C | 0.83 | -0.31 | 4.05 | WFc5PI-2 |  |
| T3 | AX-181494082<sup>d</sup> | 4 | 146.92 | G | 0.96 | -0.56 | 14.22 | WFc4PI-1 |  |
| T3 | AX-181158344<sup>c</sup> | 3 | 16.66 | C | 0.96 | -0.55 | 14.29 | WFc3PI-1 | T2, LossT |
| T3 | AX-181465065 | 4 | 1327.65 | G | 0.31 | -0.20 | 1.65 |  |  |
| T3 | AX-416776564 | 1L | 1651.35 | G | 0.38 | -0.18 | 0.99 |  |  |
| T3 | AX-416775062<sup>e</sup> | 2 | 694.64 | C | 0.45 | -0.16 | 4.55 |  |  |
| C3 | AX-416771238<sup>i</sup> | 1L | 432.98 | C | 0.15 | -0.41 | 12.12 |  |  |
| C3 | AX-416724438<sup>h</sup> | 1S | 483.98 | G | 0.80 | -0.37 | 11.38 | WFc1SPI-1 |  |
| C3 | AX-416749547 | 5 | 1285.21 | T | 0.71 | -0.27 | 5.27 | WFc5PI-3 |  |
| C3 | AX-416755845<sup>j</sup> | 1L | 1584.98 | C | 0.24 | -0.25 | 4.40 |  |  |
| LossT | AX-416775062<sup>e</sup> | 2 | 694.64 | C | 0.45 | -0.97 | 5.03 |  |  |
| LossT | AX-416799928<sup>g</sup> | 4 | 1413.44 | C | 0.88 | -1.12 | 5.43 |  |  |
| LossT | AX-416774176<sup>a</sup> | 5 | 693.09 | C | 0.94 | -1.71 | 15.88 | WFc5PI-1 |  |
| LossT | AX-416789006<sup>f</sup> | 6 | 1134.78 | C | 0.38 | -0.73 | 3.21 | WFc6PI-1 |  |
| LossC | AX-416771238<sup>i</sup> | 1L | 432.98 | C | 0.15 | -1.49 | 11.62 |  |  |
| LossC | AX-416755845<sup>j</sup> | 1L | 1584.98 | C | 0.24 | -1.09 | 5.95 |  |  |
| LossC | AX-416724438<sup>h</sup> | 1S | 483.98 | G | 0.80 | -1.14 | 6.05 | WFc1SPI-1 |  |
| LossC | AX-416730339 | 1L | 950.93 | C | 0.72 | -0.83 | 2.61 |  |  |
| LossC | AX-181493642 | 1S | 1479.49 | C | 0.11 | -1.08 | 5.81 |  |  |
| LossC | AX-416776778 | 4 | 132.45 | T | 0.72 | -0.77 | 2.95 | WFc4PI-1 |  |
| LossTC | AX-416766819<sup>b</sup> | 4 | 1523.15 | T | 0.12 | -3.12 | 19.42 | WFc4PI-2 |  |
| LossTC | AX-416775062<sup>e</sup> | 2 | 694.64 | C | 0.45 | -1.55 | 9.40 |  |  |
| DtS | AX-416746442 | 5 | 1283.70 | A | 0.72 | 3.76 | 13.80 | WFc5PI-3 |  |
| DtS | AX-416812141 | 5 | 1299.46 | T | 0.89 | 4.62 | 10.18 | WFc5PI-3 |  |
| DtS | AX-416787382 | 1S | 464.40 | T | 0.69 | 2.96 | 3.28 | WFc1SPI-1 |  |
| DtS | AX-181158344<sup>c</sup> | 3 | 16.66 | C | 0.96 | 6.14 | 12.98 | WFc3PI-1 | T2, LossT |
| DtS | AX-416809101 | 5 | 387.85 | A | 0.91 | 4.36 | 5.90 |  |  |
| DtS | AX-416789952 | 2 | 749.96 | G | 0.94 | 4.73 | 6.23 |  |  |
| T-REG | AX-181442798 | 6 | 1130.96 | C | 0.85 | 0.24 | 10.48 | WFc6PI-1 |  |
| T-REG | AX-181485355 | 6 | 759.84 | T | 0.22 | 0.22 | 8.18 |  |  |
| T-REG | AX-181165879 | 5 | 1248.26 | G | 0.22 | 0.19 | 6.41 | WFc5PI-2 |  |

Chr, chromosome; Pos, position in Mb
ben-Allele, beneficial Allele; ben-AF, allele frequency of ben-Allele; ben-Effect, ben-Allele effect
PVE, phenotypic variance explained; plQTL, pleiotropic QTL ID
MTA in LF, marker-trait association in GWAS for late-frost tolerance
a-j, denotes identical SNP marker

### GWAS on LFT

Eighteen traits were submitted to GWAS and 75 MTAs, consisting of 62 unique markers, were identified with at least two MTAs per trait (**Fig. 2**; **Table 5**; **Supplementary Figure S.R.3-4**). Nine markers showed significant MTAs with up to four different traits; mainly symptom FRS-traits, such as AX-181489050 (T1, T2, T3, LossT) and AX-416808662 (T1, C2, LossC, LossTC). We found the most MTAs for DtS (n = 9), PH1 (n = 8), LN1 (n = 7), and C1 (n = 6). PVE ranged from 1.21% (DtS) to 40.70% (T3) at the single-marker level with an average PVE of 10.06%. Overall, the trait-wise PVE ranged from 19.23% (deltaPH) to 61.60% (C1; **Table 6**). In 34 of the 62 unique markers, the major allele (avg. MAF ≈ 0.20) caused trait value improvement.

**Table 5:**
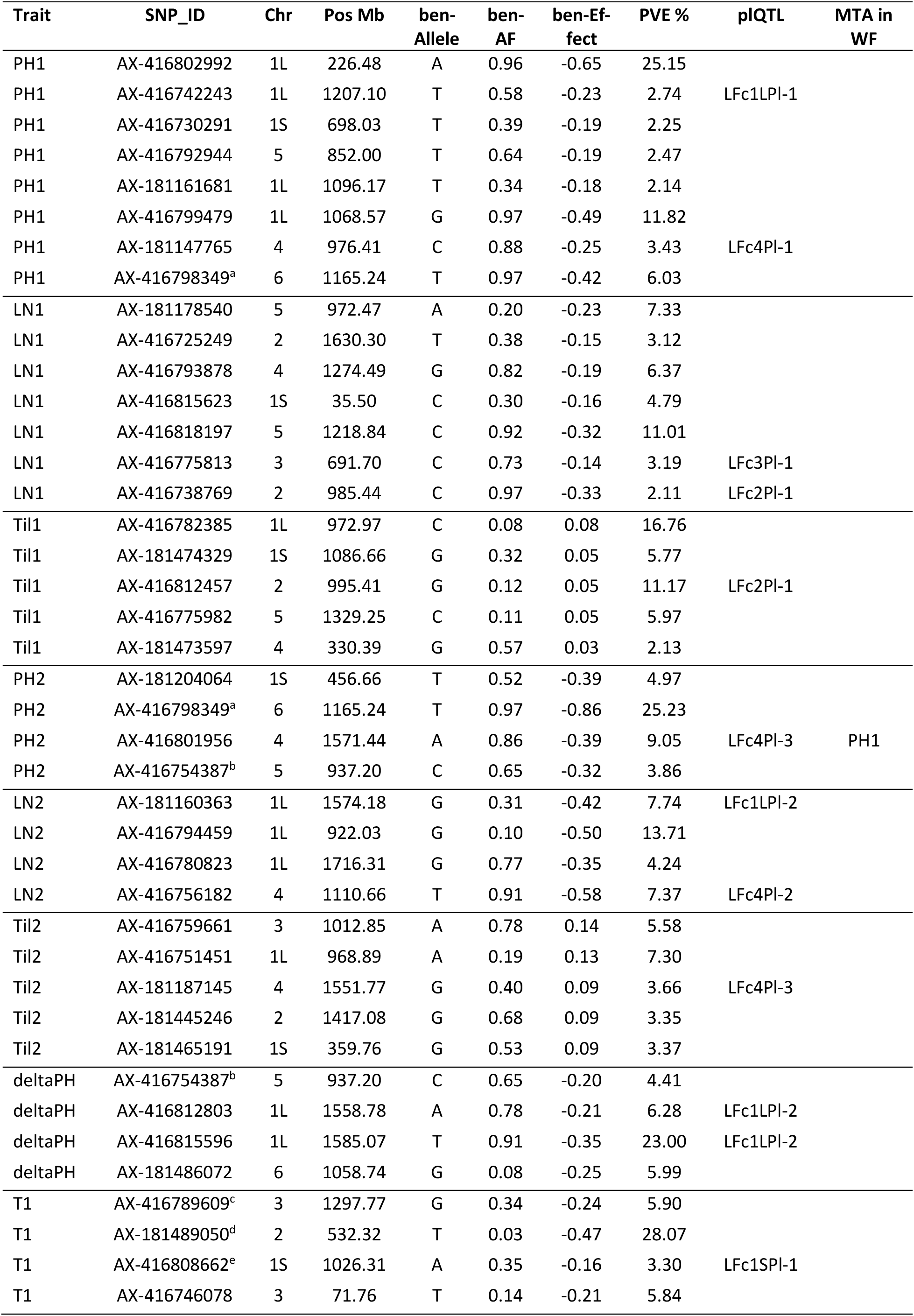
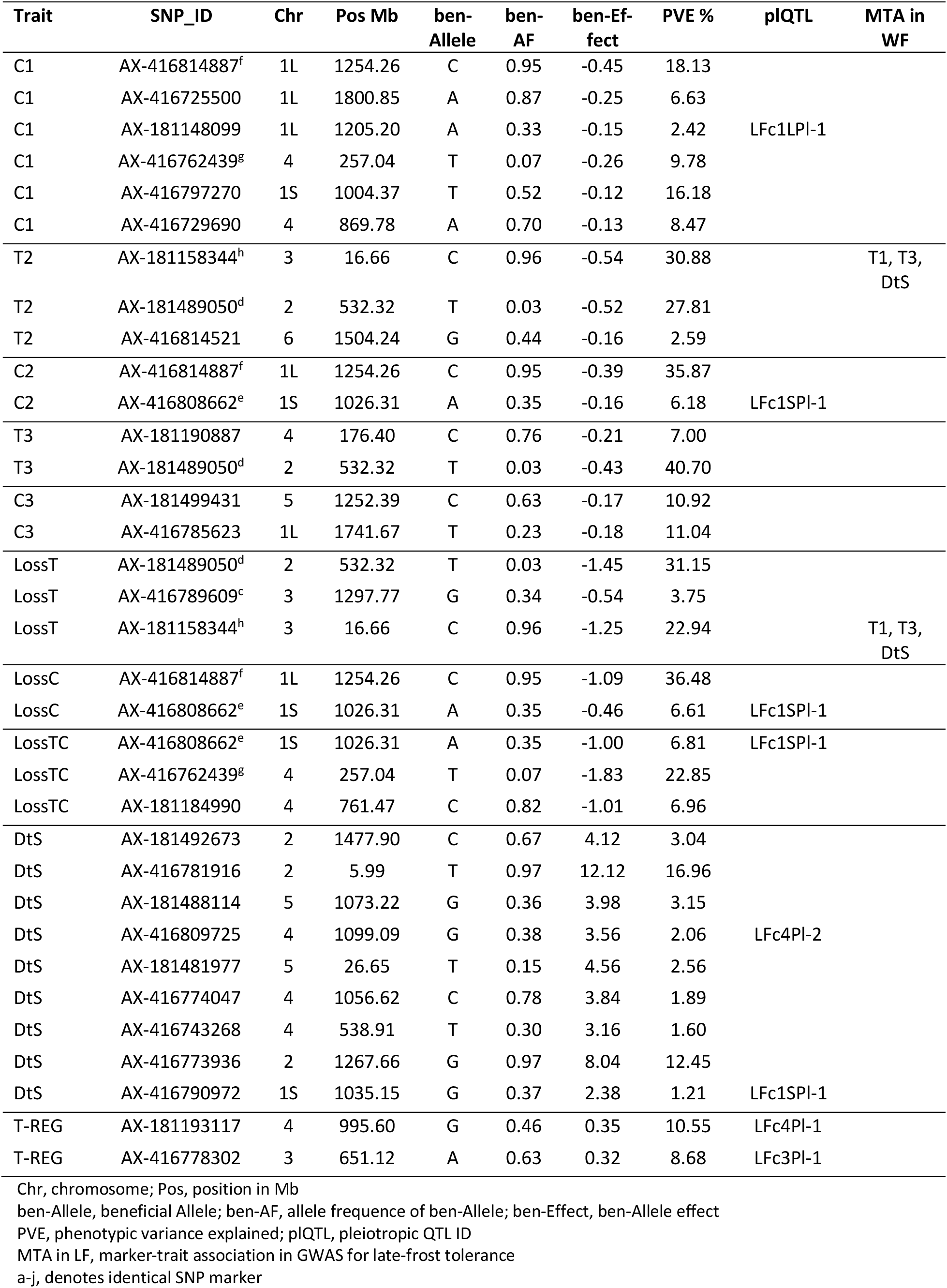
Catalog of significant marker-trait associations from genome-wide association study for late-frost tolerance. In descending order per trait according to p-value.

**Table 6:** Correlation of marker scores to phenotypic values and explained phenotypic variance by markers and marker scores within the A-set for both winter-frost and late-frost traits.

| Trait | Winter-frost |  |  | Late-frost |  |  |
| --- | --- | --- | --- | --- | --- | --- |
|  | r* | R<sup>2</sup> % | PVE % | r* | R<sup>2</sup> % | PVE % |
| PH1 | 0.50 | 24.96 | 29.67 | 0.70 | 49.56 | 56.04 |
| LN1 | 0.54 | 29.20 | 39.16 | 0.66 | 43.15 | 37.92 |
| Til1 | - | - | - | 0.59 | 34.95 | 41.81 |
| PH2 | - | - | - | 0.55 | 30.20 | 43.10 |
| LN2 | - | - | - | 0.58 | 33.35 | 33.05 |
| Til2 | - | - | - | 0.60 | 35.43 | 23.26 |
| deltaPH | - | - | - | 0.55 | 29.76 | 39.68 |
| T1 | 0.59 | 35.05 | 56.11 | 0.58 | 34.08 | 43.10 |
| C1 | 0.68 | 45.83 | 45.13 | 0.65 | 41.80 | 61.60 |
| T2 | 0.51 | 26.34 | 30.49 | 0.45 | 20.06 | 61.27 |
| C2 | 0.57 | 32.81 | 49.01 | 0.45 | 19.93 | 42.05 |
| T3 | 0.73 | 52.84 | 58.12 | 0.45 | 20.68 | 47.71 |
| C3 | 0.51 | 26.46 | 33.16 | 0.43 | 18.81 | 21.96 |
| LossT | 0.51 | 26.28 | 29.55 | 0.48 | 23.33 | 57.84 |
| LossC | 0.54 | 29.42 | 35.00 | 0.45 | 20.51 | 43.10 |
| LossTC | 0.42 | 17.41 | 28.82 | 0.45 | 20.61 | 36.62 |
| DtS | 0.70 | 49.35 | 52.37 | 0.75 | 56.88 | 44.92 |
| T-REG | 0.50 | 24.61 | 25.07 | 0.43 | 18.62 | 19.23 |
| Correlation to number of markers |  | 0.86 | 0.62 |  | 0.98 | 0.30 |

r, Pearson correlation coefficient (prediction ability)
r\*, r significant at p-value ≤0.001
R<sup>2</sup>, squared correlation of marker score to phenotypic values
PVE, phenotypic variance explained by all markers per trait

### Pleiotropic QTL Identification

We identified 18 candidate regions of putative pleiotropic QTLs (plQTL) both within and across treatments (**Table 7**; **Supplementary Table S.R.6**). On single-marker level, four markers were considered to address plQTL directly, as they were each significantly associated with at least two traits of different trait categories. For example, marker AX-416774176 on chr5 associated with multiple WF symptom FRS-traits as plQTL WFc5Pl-1. Similarly, marker AX-181158344 on chr3 affects multiple FRS-traits as plQTL FTc3Pl-1 across treatments.

**Table 7:** Identified putative QTL and pleiotropic QTL (plQTL) with and across winter-frost (WF) and late-frost (LF) treatments. Each marker was counted only once as either associated with a QTL or a plQTL.

| Trait type | Traits | QTL |  |  | n QTL | Pleiotropic QTL |  |  | n pQTL |
| --- | --- | --- | --- | --- | --- | --- | --- | --- | --- |
|  |  | WF | LF | WF & LF |  | WF | LF | WF & LF |  |
| non-FRS-traits | PH1/2, LN1/2, Til1/2, deltaPH | 3 | 17 | 1 | 21 | 0 | 1 | 1 | 2 |
| mix FRS-traits | Symptoms and survival | - | - | - | - | 2 | 0 | 2 | 4 |
| Symptoms | T loss | 4 | 5 | 0 | 9 | - | - | - | - |
|  | C loss | 3 | 2 | 0 | 7 | - | - | - | - |
|  | T and C loss | - | - | - | - | 1 | 0 | 1 | 2 |
| Survival | DtS | 2 | 7 | 0 | 9 | - | - | - | - |
|  | T-REG | 1 | 0 | 0 | 1 | - | - | - | - |
|  | DtS and T-REG | - | - | - | - | 0 | 0 | 0 | 0 |
| unspecific* | non-FRS- and FRS-traits | - | - | - | - | 0 | 1 | 9 | 10 |
| Total |  | 15 | 31 | 1 | 47 | 3 | 2 | 13 | 18 |

n QTL, total number of QTL
n pQTL, total number of pleiotropic QTL
\*, pQTL affects both non-FRS- and FRS-traits

At treatment level, we found eight treatment-specific plQTLs each (**Table 7**). In WF, five plQTLs affected FRS-traits and three plQTL only affected symptom FRS-traits, such as WFc1SPl-1 (DtS and color loss) on chr1S or WFc4Pl-1 (LossC, T1, and T3) on chr4, respectively. In LF, in contrast, only LFc1SPl-1 on chr1S affected exclusively FRS-traits (DtS, C1, T1, C2, LossC, and LossTC) and three plQTL affect only non-FRS-traits, while most plQTL (n = 4) affected both non-and FRS-traits. Traits affected by the same putative plQTL were mostly highly correlated, such as LossC, T1, and T3 in WFC4Pl-1 on chr4.

At the cross-treatment level, most treatment-specific plQTLs became treatment-unspecific due to physical vicinity to MTAs discovered in the other treatment (**Supplementary Table S.R.6**). For example, WFc1SPl-1 and the marker AX-181204064 for PH2 (LF) were combined to FTc1SPl-1 on chr1S. Consequently, only three WF-specific and two LF-specific plQTLs remained, while 13 treatment-unspecific plQTL were identified. Treatment-unspecific plQTL were associated with up to four different traits of maximal two trait categories, such FTc5Pl-1 (**Fig. 3**). Interestingly, all plQTL included at least one treatment-specific plQTL, except FTc6Pl-1 on chr6.

**Fig. 3:**
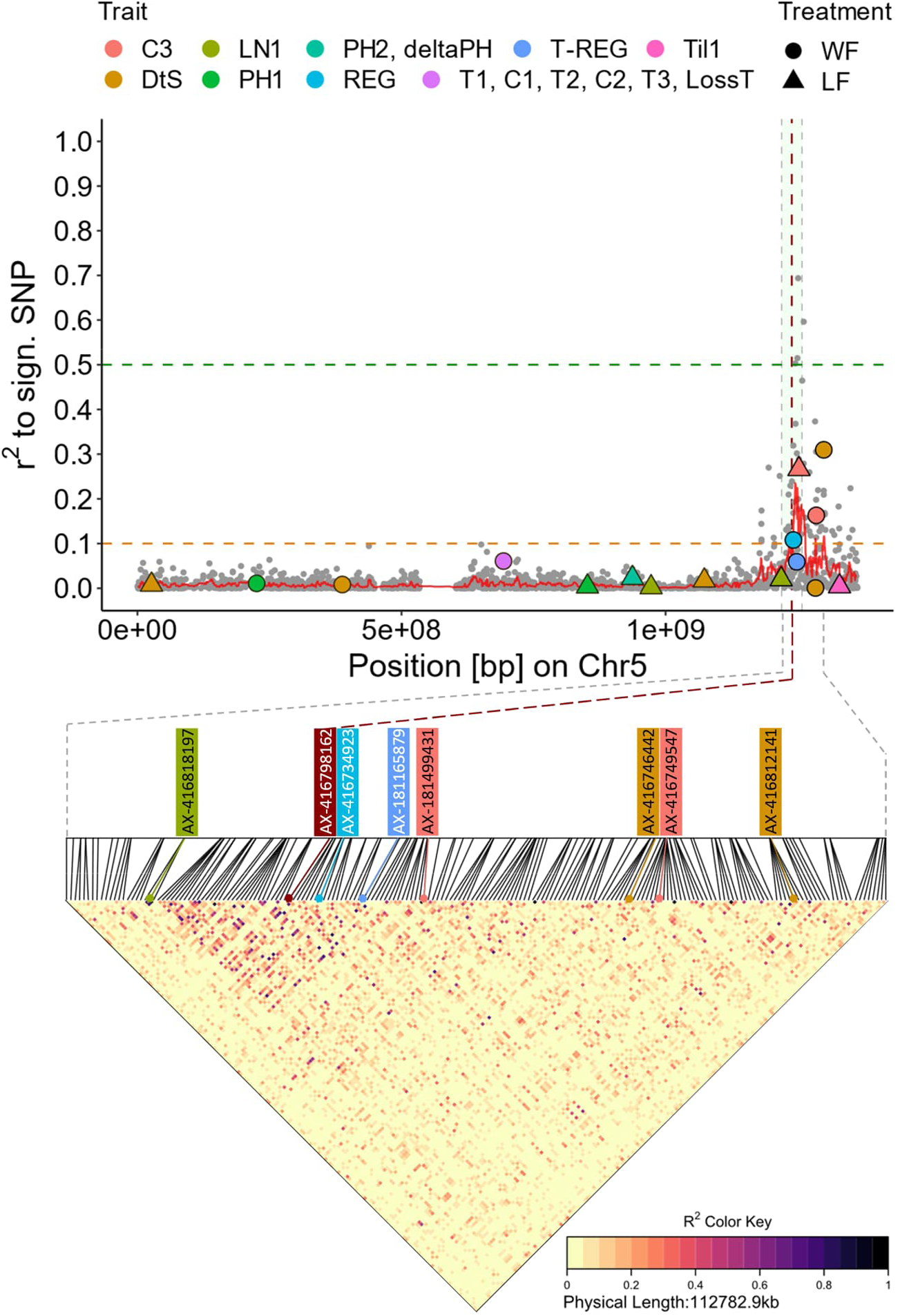
Local LD (r²) analysis among significant associated markers (colored) identified on chromosome 5 across all traits and both the winter-frost (circle) and late-frost (triangle) treatment. The genomic region of putative pleiotropic QTL FTc5Pl-1 (**Fig. R.2**) with pairwise SNP LD is shown as LDheatmap below. Gray dots: non-significant SNPs; colored dots and triangles, significant SNPs; vertical red dashed line, position of marker AX-416798162 associated with T3; light-green shaded area, indicates LD decay region to each side of AX-416798162; red line, smoothing curve of local LD in 1000er bins.

Analysis of pairwise marker LD within each plQTL yielded often substantially higher LD than expected at the given physical distance (**Fig. 3-4**). Within LFc1SPl-1, for instance, the markers AX-416790972 (DtS) and AX-416808662 (T1, C2, LossC, and LossTC) were in high LD (r² = 0.16). Within FTc5Pl-1, pair-wise LD was even up to of r² = 0.31 between the markers AX-416812141 (DtS; WF) and AX-416798162 (T3; WF; **Fig. 3**). However, some markers within a plQTL showed little to no LD such as in FTc1SPl-1 or FTc1LPl-1 with a pairwise LD of r² ≤0.09.

**Fig. 4:**
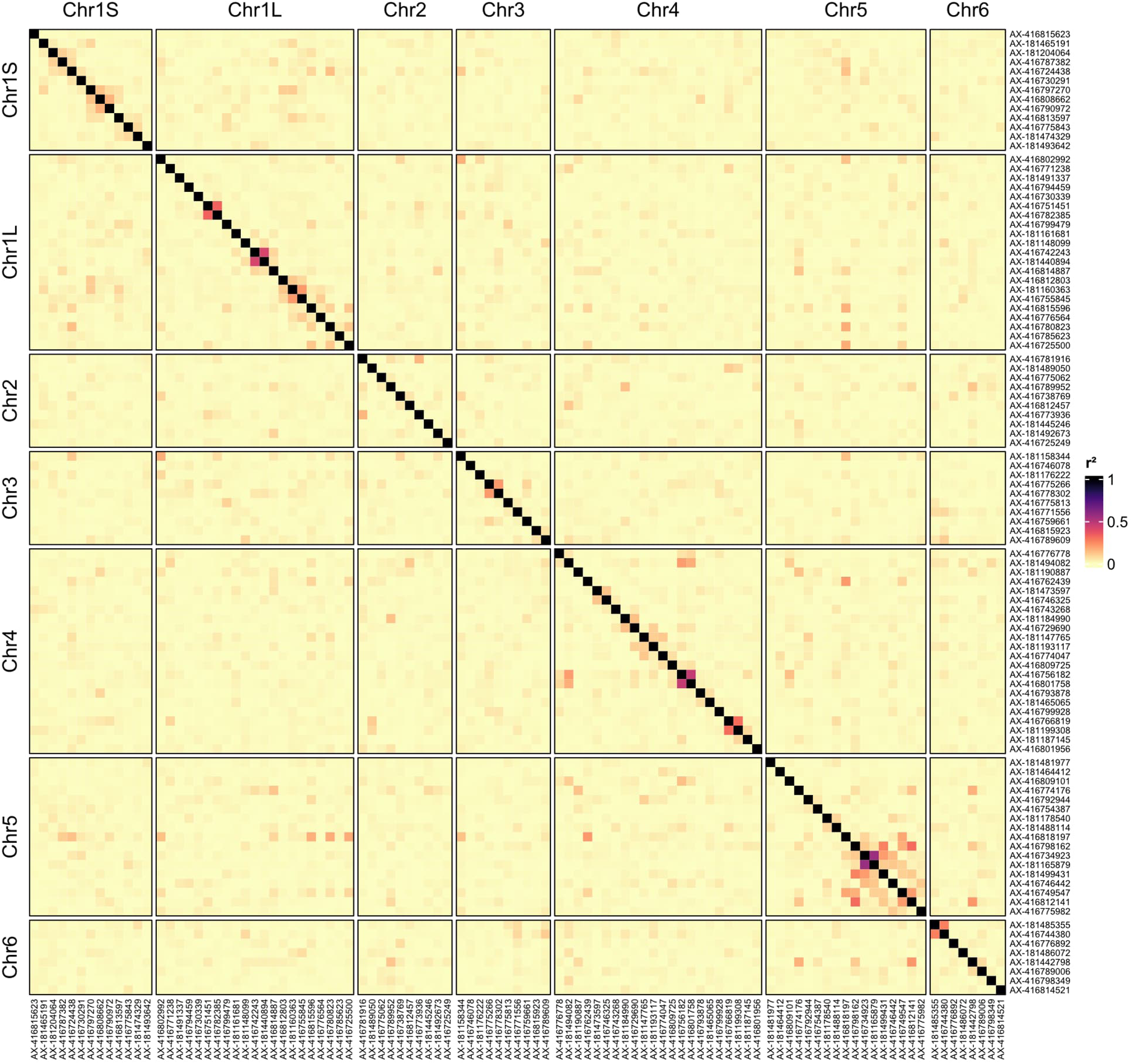
LD heatmap that displays pairwise LD (r²) across all significant markers on all chromosomes identified by GWAS for winter-frost and late-frost tolerance.

### Marker Score-based Prediction

To evaluate the predictive power of identified markers, we calculated trait-specific marker scores per A-set line and estimated prediction ability via correlation to the respective phenotypic values (**Fig. 5**; **Table 6**). We found medium to high prediction abilities (0.42-0.75; p ≤0.05) for all traits in both treatments. For the WF traits, the marker scores could explain on average 31.39% of the phenotypic variance, with a range of 17.41% (LossTC) ≤ R² ≤ 52.84% (T3). A similar range and average of R² were observed for the LF traits: 17.88% (REG) to 56.88% (DtS), on average 29.98%. R² was highly correlated with the number of contributing markers (0.86/0.98 in WF/LF).

**Fig. 5:**
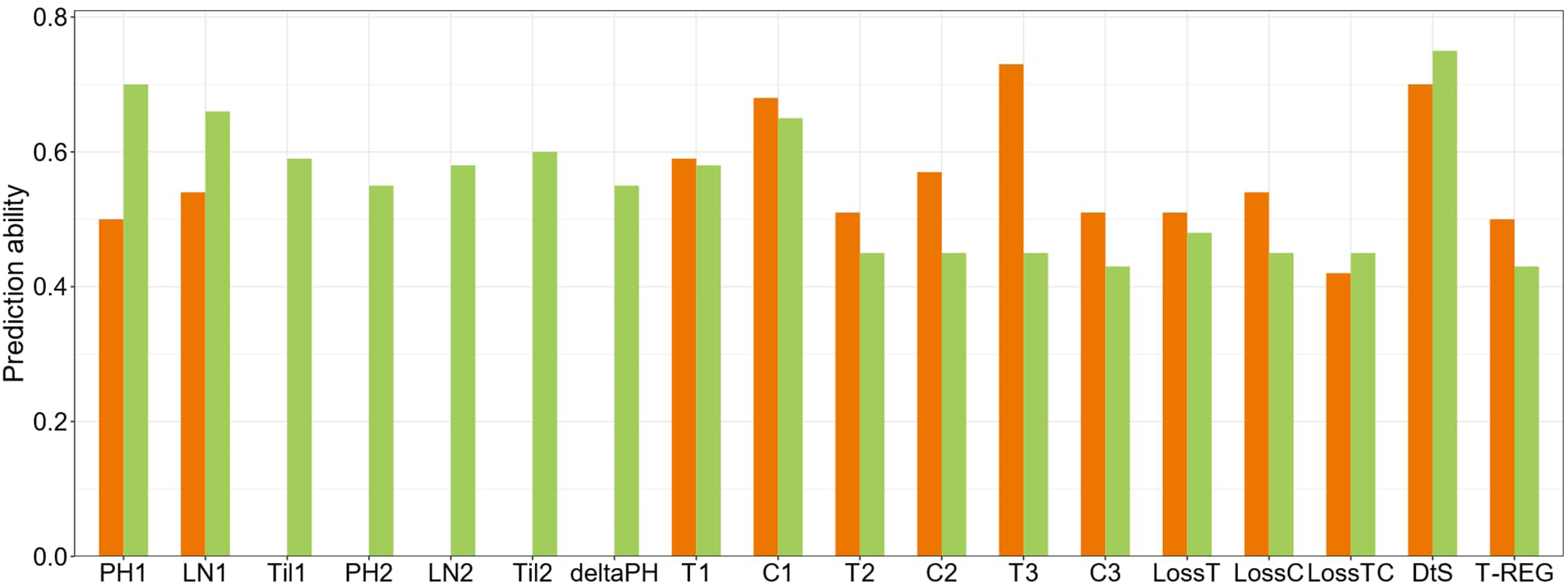
Prediction abilities of marker scores for traits in winter-frost (orange) and late-frost (green) treatment. Marker scores were calculated from BLINK-estimated marker effects of all significant associated markers per trait to estimate explained phenotypic variance within the A-set. Prediction ability estimated as Pearson correlation of the marker score to the observed phenotypic value per trait.

## Discussion

### Genome-Wide Analysis Underlines Independent Genetic Architecture of Winter-Frost and Late-Frost Tolerance

Attractiveness of faba bean as a winter crop suffers from the risk of significant yield losses by winter kill, especially in Central and Northern Europe. To this end, breeding with substantial genetic gain for winter hardiness in winter faba bean is essential. However, winter hardiness is a complex component trait, affected by numerous abiotic and biotic stresses (Junttila 1996). Thus, breeding progress is slow (Michel et al. 2019) and lack of frequent, well-differentiating cold winters for reliable selection hampers it even further. Therefore, artificial frost tests have become the standard approach to screening for freezing tolerance (FT), the major component of winter hardiness (Duc et al. 2011). Previous studies have already detected a few large-effect QTLs associated with FT under controlled conditions in faba bean, while leaving the greater proportion of phenotypic variance unexplained (Arbaoui et al. 2008; Ali et al. 2016; Sallam et al. 2016; Carillo-Perdomo et al. 2022; Windhorst et al. 2023). In accordance with these findings, our GWAS for winter-frost tolerance yielded seven major QTLs (PVE ≥10%) across all FRS-traits and 30 MTAs in total.

In fact, the major QTL for turgor loss on chr2, marker AX-416775062, co-localized (distance: approx. 3 cM) with the marker Vf_Mt3g086600 associated with LossTC and DtS identified by Ali et al. (2016). Interestingly, we mapped a QTL for DtS (WF) within 50 Mb distance to the turgor loss QTL, rendering the whole genomic region as highly interesting for candidate gene search. The second marker (Vf_Mt5g046030) from their study, associated with LossTC, did not co-localize with any QTL detected on chr1L. However, local LD pattern supports the assumption that this marker pinpoints the QTL for color loss (C1) addressed by marker AX-181491337 (PVE: 7.28%). Carillo-Perdomo et al. (2022) reported a major FT QTL on chr1L that co-localized with a previously reported QTL for FT and fatty acid composition (Sallam et al. 2016). The markers flanking this QTL, however, span an approx. 600 Mb region (pos: 935-1,548 Mb), within which GWAS now yielded three MTAs for color loss, including two detected in LF treatment. LD among the three MTAs suggests independent QTLs rather than a single treatment-unspecific major QTL for color loss. Therefore, precise co-localization to the FT QTL reported by Carillo-Perdomo et al. (2022) was not verifiable. However, the A-set originated from the winter hardy GWBP and possesses a varying (**Table 3**) but generally high FT level, i.e. winter hardiness. Accordingly, it was not expected to detect key FT loci differentiating spring-type from winter-type, as it was the case in Carillo-Perdomo et al. (2022).

In addition, our GWAS results for winter-frost tolerance provide further insights into the highly quantitative nature of this trait since we identified 2-8 MTAs per FRS-trait, that explained up to 58.12% of observed phenotypic variance. Having detected both QTL for each FRS-trait but also plQTL, that affect multiple FRS-traits, such as WFc1SPl-1 for color loss and DtS on chr1S (**Table 7**; **Supplementary Table S.R.6**), our results further indicate multiple pathways being involved in winter-frost tolerance (Xin and Browse 2000). For example, protection against cell damage in leaves and stems (color and turgor loss) involves not only several QTL commonly detected at all frost stress levels (T1/C1, T2/C2, T3/C3), but also few QTL specifically detected only after the first frost stress (T1/C1). This could indicate a distinct mechanism for protection against cell damage upon initial frost stress in cold acclimated plants.

While cold acclimation provides freezing tolerance in winter (Herzog 1989; Thomashow 1999), plants deharden and become sensitive to freezing in spring as temperatures and day length increase (Junttila 1996). Therefore, late-frost in spring poses a contrasting challenge to faba bean plants. However, assessment of late-frost tolerance in the field is even more difficult and screening in controlled conditions more laborious than for winter-frost tolerance. It is therefore not surprising that no association or QTL mapping study has been reported for this trait in winter faba bean. In this study, we found great genetic variance for late-frost tolerance in the A-set suggesting good potential for trait improvement. In fact, GWAS identified 25 MTAs across all FRS-traits of which ten were associated with major effect QTL (**Tables 4-5**). In contrast to the WF GWAS results, we found only one plQTL, LFc1SPl-1 on chr1S, associated with multiple LF FRS-traits: turgor loss, color loss, and DtS (**Supplementary Table S.R.6**). Considering the less strong phenotypic and genetic correlations between turgor and color loss than in the WF treatment, reversible turgor loss vs irreparable cell damage (color loss) in leaves and stems appears to involve fewer interdependent pathways than in hardened plants under winter-frost stress. Long-term survival of severe late-frost, however, involves an entirely different set of QTLs; similar to WF treatment.

Considering the different physiological state of plants in spring compared to winter, we hypothesized an independent quantitative genetic control of late-frost vs winter-frost tolerance. Indeed, we not only observed low phenotypic correlations (r ≤0.50; **Supplementary Table S.R.3**) of the same FRS-traits between WF and LF treatment but detected 18 (WF) and 33 (LF) treatment-specific QTL (**Fig. 2-3**; **Tables 3-4** and **7**; **Supplementary Table S.R.6**). Contrastingly and as expected, PH1 and LN1, unaffected by frost treatments, correlated strongly (r >0.71; **Supplementary Table S.R.3**) between the treatments; hence the common MTAs for PH1 (PVE: 17.72%, WF) and PH2 (PVE: 9.05%, LF) on chr4. Moreover, major QTLs for turgor loss, color loss, DtS, and regrowth detected per treatment showed almost no association across treatments. In addition, only five of the 13 treatment-unspecific plQTLs affect FRStraits in both treatments. For example, the major plQTL WFc5Pl-3 on chr5 explained the greatest proportion of DtS in response to winter-frost, while most major QTL for DtS after late-frost are located on chr2. Altogether, our results indicate treatment-specific effects of putative QTLs, which strongly contribute to rank changes in lines FT across treatments. However, for both turgor loss and color loss we found at least one treatment-unspecific plQTL (FTc3PL-1, FTc5Pl-1), suggesting a common base in genetic control as well (**Supplementary Table S.R.6**). Especially, FTc3Pl-1 with its major effect on turgor loss (WF and LF) and DtS (WF) as well as the huge plQTL FTc5Pl-1 on chr5 (**Fig. 3**) seems to play a pivotal role in winter faba bean freezing tolerance, as they affect not only symptom expression in leaves and stem but also the long-term survival in both winter-frost and late-frost stress.

### Usability for MAS

The detection of major effect QTLs for all traits (PVE ≤40.70%; **Tables 3-4**) suggests great potential for MAS. This was supported by the medium to high, significant prediction abilities for marker scores obtained for all FRS-traits in the A-set (0.42≤ r ≤0.75; avg. 0.53; **Fig. 5**). In other words, the marker score could explain >25% (R²) of the phenotypic variance in many LF traits and in almost all WF traits. Especially for C1 and T3 in WF, T1 and C1 in LF, and DtS in both treatments, we captured ≥50% of the phenotypic variance in the A-set. The low prediction ability obtained for LossTC (WF) and T-REG (WF and LF) could be explained by the limited number of identified MTAs, as R² was highly correlated to the number of markers; 0.86 (WF) and 0.98 (LF; **Table 6**).

Although these findings seem promising for successful MAS-based genetic gain for freezing tolerance, final validation of markers is required, as prediction ability was only evaluated within the A-set without a (cross-)validation approach. Unfortunately, reproducibility of GWAS-based MTAs is often limited by the limitations of the GWAS design itself (see below). In addition, a breeder would face two major hurdles when transferring these markers into a winter faba bean breeding program; (1) the marker and the QTL must segregate and (2) the LD and linkage phase orientation between marker and QTL must be conserved in the breeding material. Whether these prerequisites are met depends on the breeding material, its breeding history, its relatedness to the GWAS panel (Daetwyler et al. 2013; Hickey et al. 2014) as well as the extent of LD (Windhausen et al. 2012) and both the marker and QTL allele frequency distribution (Hayes 2013).

We have therefore developed a two-step validation procedure that addresses the issue of marker transferability. To facilitate powerful hard-validation, we phenotyped an independent set of n = 64 winter faba bean inbred lines with varying levels relatedness to the A-set and evaluated both the MTAs and marker effect sizes. The results of this second part of the current study are in preparation and will be presented in the final publication.

### Limitations of the GWAS

GWAS relies on the LD or linkage between SNP markers and QTL and discovery of MTAs depends mainly on (1) accurate phenotypic data, (2) number and genome coverage of markers, (3) LD, (4) number and effect sizes of QTL, and (8) the GWAS panel size; in short, the power of detection (Bush and Moore 2012; Spencer et al 2009; Wang and Xu 2019).

The discovery of MTAs in this study was potentially impaired by the trait’s genetic architecture and h², extent of LD, and size and composition of the A-set itself. The rather low average pairwise LD of adjacent SNPs (r² = 0.15) and the comparable fast LD decay in the A-set (Skovbjerg et al. 2023) should have favored the GWAS resolution. However, given the observed SNP-to-gene coverage (approx. 1 SNP/2 genes; **Supplementary Table S.R.4**) and evenly distributed recombination sites along the whole chromosome in the faba bean genome (Jayakodi et al. 2023), small-effect QTL still would have been difficult to detect due to low LD, especially at low MAF (<0.10).

In addition, the size of the A-set (max. n = 185) could have had a substantial impact on QTL discovery. At small sample size (<200), discovery of minor effect QTL is difficult as power of detection is rather low (Bradbury et al. 2011; Land and Thompson 1989). In fact, at the given sample size and average pairwise LD between SNPs, power of detecting QTL with 2.5-5% PVE could be assumed approx. 0.10-0.40 according to simulations by Hayes (2013) and was even further compromised by low MAF (Wang and Xu 2019). Inevitably, a minor SNP allele (MAF <0.10; equal to <18 lines) was present in only a few lines. Consequently, BLINK-based PVE by a single marker was most likely overestimated due to panel size (Melchinger et al. 1998), a phenomenon known as Beavis effect (Beavis 1994; Sahana et al. 2023; Xu 2003), and due to the non-representative genetic background and/or extreme phenotypes in the few lines carrying the minor allele (Shana et al 2023). This bias can be inferred from the exponential decay of PVE by increasing MAF (**Supplementary Fig. S.5-6**) observed for all markers. In fact, the total PVE per trait as provided by BLINK was up to 41.21% higher than the respective marker score-based R² (**Table 6**). For example, two of the three markers for LossT (LF) accounted for 22.94% and 31.15% PVE at very low MAF ≤0.04, whereas the marker score-based R² using all three markers was only 23.33%. The proportion of markers of MAF <0.10 was especially high among LF traits (15/63 markers). The effect sizes of these markers could have been drastically overestimated, if not false positive, despite the FDR threshold of (p <0.05). These assumptions warrant further research during the two-step validation.

As stated above, this very winter faba bean material was suitable to detect some and predominantly larger-effect QTLs. Considering the high h² of most traits, the missing 50-75% of phenotypic variance not explained by the identified markers (**==Fig. 6**; **Table 6**) indicates that several to very many QTL with small to minimal effects per trait remain undiscovered. Their detection would require an enlarged GWAS panel (Hayes 2013) and further phenotyping under climate chamber conditions, i.e. trade-off in sample size vs number and duration of experiments. However, success in detecting additional minor QTLs would not be guaranteed (Bradbury et al. 2011). Alternatively, one could cross A-set lines carrying as few beneficial alleles for as many reported FRS-trait QTLs as possible and conduct an F_2_-based QTL-mapping. Detection of formerly rare allele QTLs would be easier (Bernardo 2020), provided a sufficiently large F_2_ population to narrow down confidence intervals (Würschum 2012). However, QTLs contributing only small additive genetic variance (<5%) to the overall phenotypic value promise only little genetic gain by MAS. Instead, one could “simply” select phenotypically for high freezing tolerance in such an F_2_ and following generations, to foster enrichment and fixation of beneficial alleles at yet unknown minor QTLs (phenotypic background selection). Inbred lines derived from this selection procedure could elevate the freezing tolerance level in the breeding material upon re-introgression provided a well-established MAS foreground selection for major QTLs (“tolerance allele pyramiding”; Collard et al. 2007).

### Candidate Gene Search Approach

The identification of candidate genes affecting freezing tolerance either in general or specifically under winter-frost or late-frost conditions, would be of great interest for molecular breeding in faba bean. Considering the average SNP density of 1.52 SNP/Mb, the LD decay of 1.32 Mb to half (r² = 0.08) of its maximum, and the approx. SNP-to-gene ratio of 1SNP/2 genes, always the neighboring genes of a marker could be assumed as most promising candidate genes. However, this candidate gene search approach would probably be oversimplified, as chromosome-wide LD decayed slower (on avg.: 17.57 Mb) and both gene and SNP distribution are not homogenous along the chromosomes. Especially for pleiotropic QTLs, that combine multiple markers over large physical genomic distances, such as the 80 Mb sized FTc5Pl-1 on chr5 (**Fig. 4**), candidate gene identification must be approached differently. Accordingly, the reported markers may be considered as physical orientation for further fine-mapping of QTLs and candidate genes rather than final target loci. Recent advancements in faba bean genomic tools, such as implementation of single primer enrichment technology (SPET) genotyping with 90K SNP (Jajakodi et al., 2023), faba bean specific Genotyping-by-Sequencing (GBS) protocols (Zhang et al. 2024), or whole-genome re-sequencing, may facilitate such study with relative ease.

## Conclusion

The here reported marker and putative QTL represent the next milestone toward unraveling the genetic architecture of FT and winter hardiness in faba bean and legumes in general. The catalog of GWAS-based MTAs, along with the new late-frost phenotypic data, provide a valuable resource for identifying key haplotypes and candidate genes. Following investigations should aim for high-resolution fine mapping within the detected genomic candidate regions to narrow down the QTL to causative or perfectly linked SNPs to facilitate successful MAS. Furthermore, incorporation of such markers as covariates into genomic prediction models could drastically improve predictive power (Michel et al. 2019). In fact, as expensive and difficult to assess highly polygenic traits, winter hardiness and freezing tolerance represent optimal targets for genomic selection. A comprehensive study that will shed first light on this matter is currently under preparation. In conclusion, the rapid developments in the field of faba bean genomics will acceleration in breeding for improved winter hardiness in the nearby future.

## Abbreviations

FT: Freezing tolerance
WF: Winter-frost
LF: Late-frost
DtS: Disposition to survive
T-REG: Transformed regrowth
MTA: Marker-trait association
PVE: Phenotypic variance explained
plQTL: Pleiotropic QTL

## Acknowledgement

A.W. acknowledges a scholarship from the Karl Eigen and Dr.h.c. Dietrich Brauer Foundation (UFOP, Union zur Förderung von Oel-und Proteinpflanzen e.V.). The authors would like to thank Ahmed Sallam for the conducted winter-frost experiments and detailed phenotyping. This research was part of the project ProFaba. ProFaba was carried out under the ERA-NET Cofund SusCrop (Grant No. 771134), being part of the Joint Programming Initiative on Agriculture, Food Security and Climate Change (FACCE-JPI). This research was funded by the Federal Ministry of Education and Research (BMBF, Grant No.: 031B0805A).

## Conflict of Interest

The authors declare no conflict of interest.

## Author Contributions

Conceptualization, W.L., A.W., S.U.A.; Methodology, A.W.; Software, A.W., C.K.S; Validation, A.W., W.L..; Formal Analysis, A.W.; Investigation, A.W..; Resources, W.L., D.M.O., D.A., S.U.A; Data Curation, A.W., C.K.S., D.A.; Writing – Original Draft, A.W.; Writing – Review & Editing, A.W., W.L.; Visualization, A.W.; Supervision, W.L., S.U.A., D.M.O.; Project Administration, W.L., S.U.A., D.M.O.; Funding Acquisition, W.L., S.U.A., D.M.O..

## Data Availability

Relevant data supporting the presented findings are provided within the paper and its Supplementary Figures and Tables.

## Supplementary Figures

**Figure S.1:**
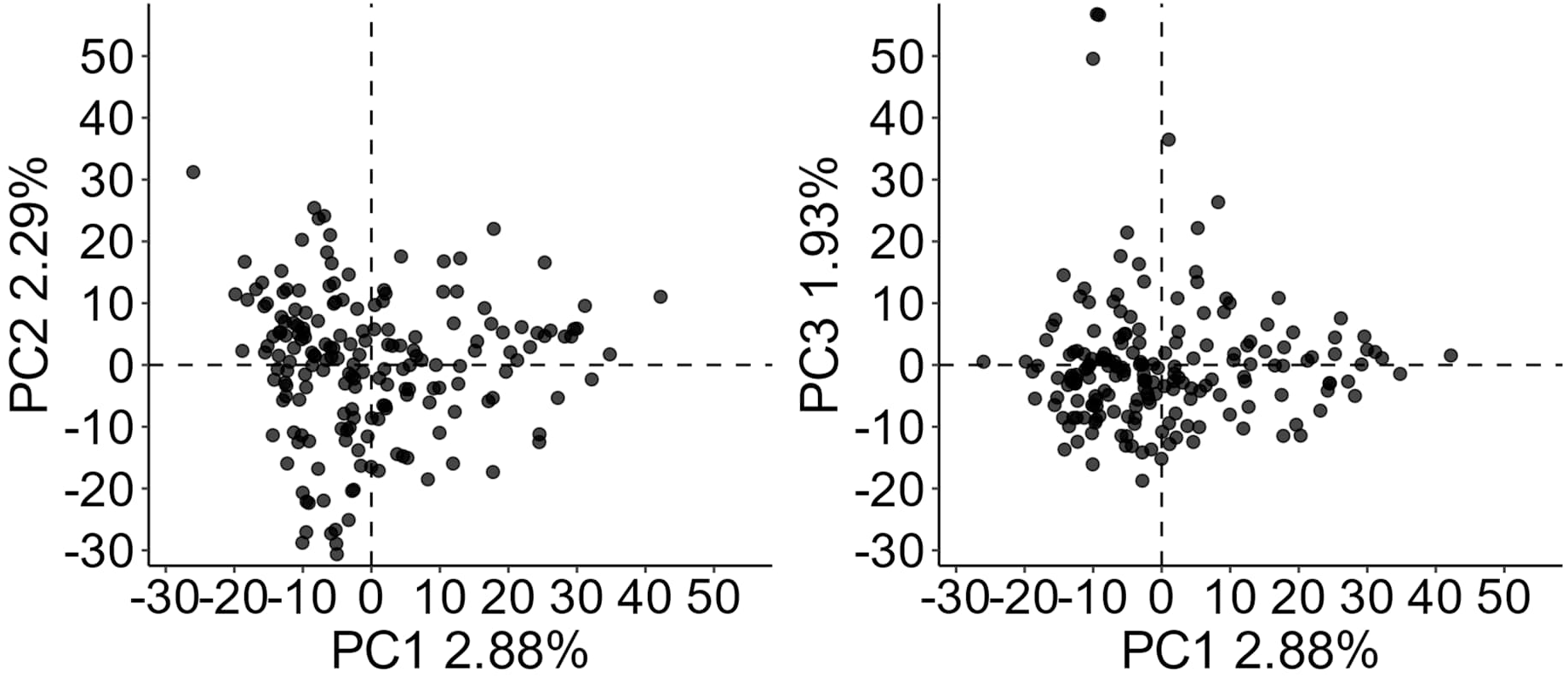
Principal component analysis of A-set (n = 185).

**Figure S.2:**
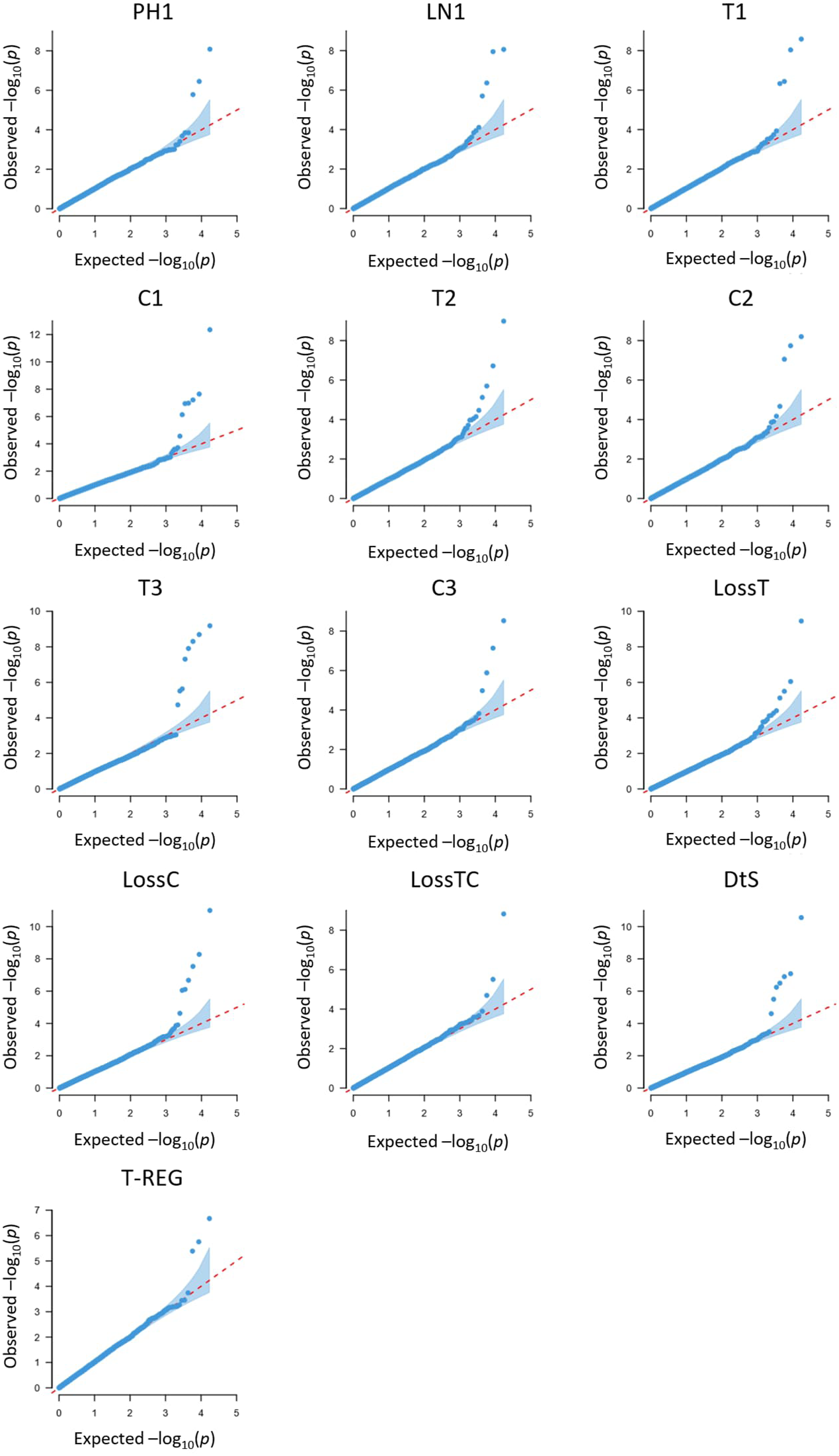
QQ-plots of 13 traits analyzed in winter-frost GWAS.

**Figure S.3:**
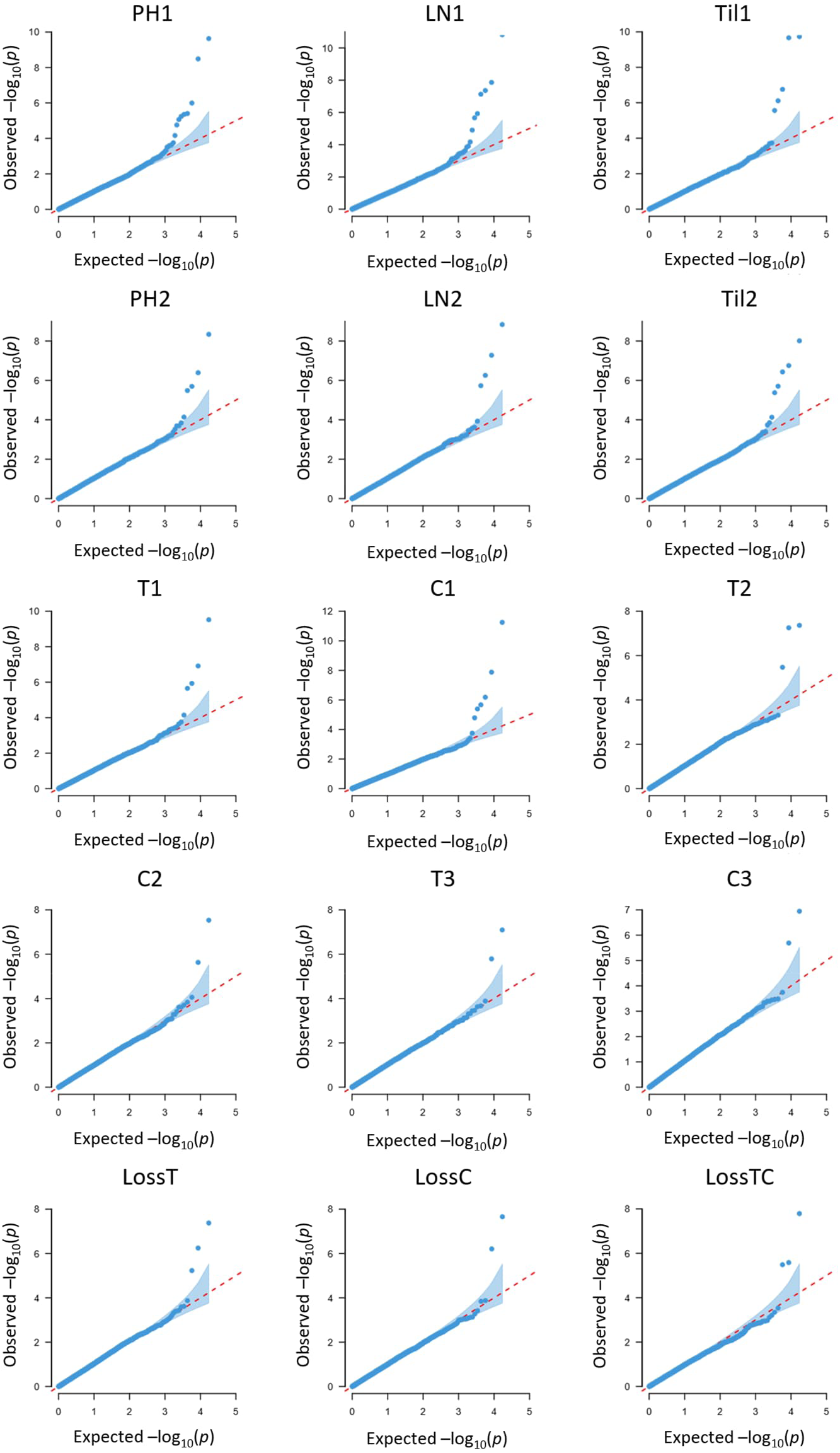
QQ plots of 18 traits analyzed in late-frost GWAS. Traits DtS, T-REG, and deltaPH displayed in **Figure S.4**.

**Figure S.4:**
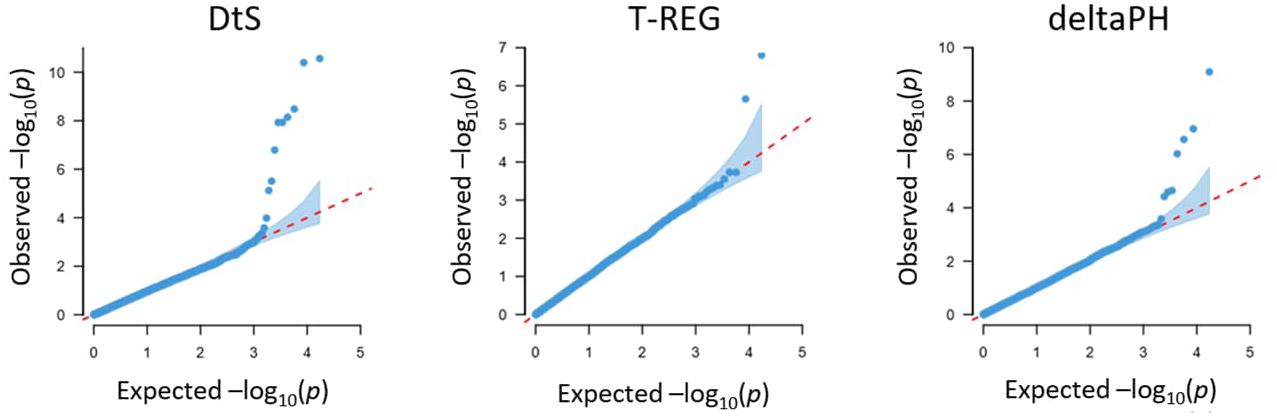
QQ plots for DtS, T-REG, and deltaPH analyzed in GWAS for late-frost tolerance.

**Figure S.5:**
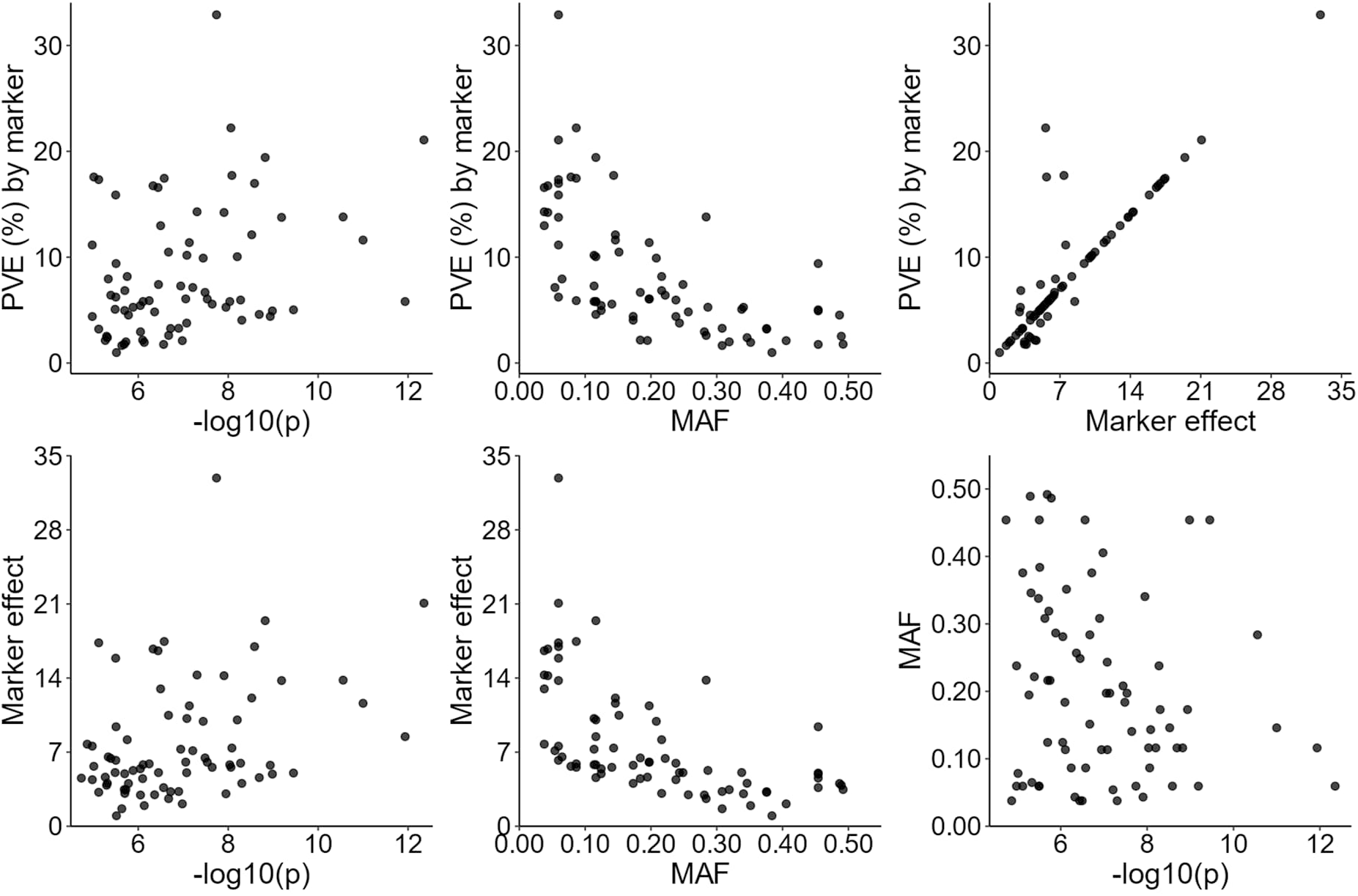
Scatter plots comparing the phenotypic variance explained (PVE), minor allele frequency (MAF), the -log(10) p-value, and the marker effect of per significant marker identified in winter-frost GWAS.

**Figure S.6:**
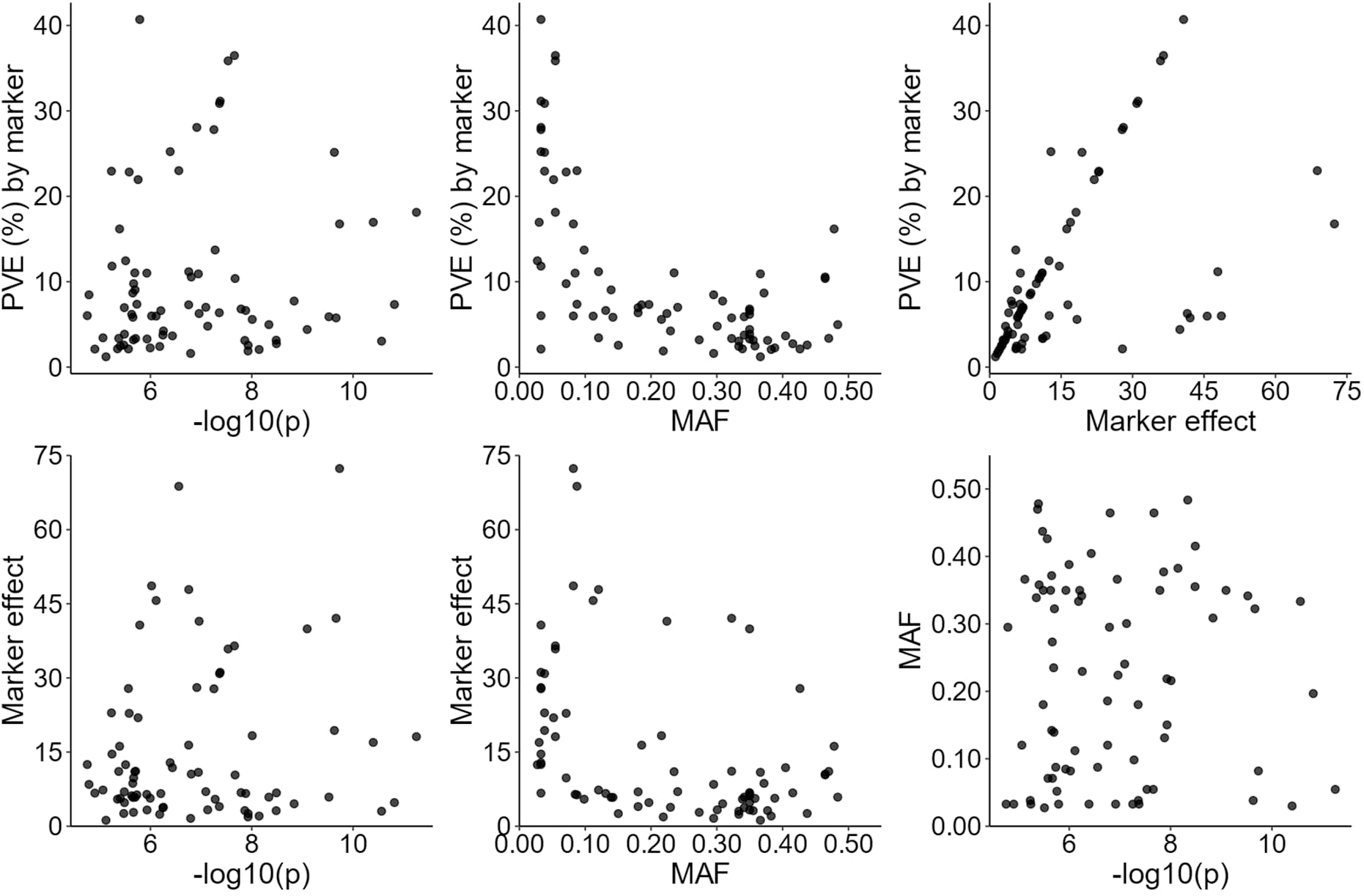
Scatter plots comparing the phenotypic variance explained (PVE), minor allele frequency (MAF), the -log(10) p-value, and the marker effect of per significant marker identified in winter-frost GWAS.

## Notes

### Competing Interest Statement

The authors have declared no competing interest.

